# Electrophysiological abnormalities associated with a *CACNA1D* variant are rescued by AAV6-Cav1.3-C-terminus gene therapy in patient-iPSC-CMs

**DOI:** 10.64898/2025.12.15.694396

**Authors:** Jean-Baptiste Reisqs, Yvonne Sleiman, Ivan Gando, Michael Cupelli, Vamsi Krishna Murthy Ginjupalli, Helene Delanoe-Ayari, William Coetzee, Yongxia S. Qu, Mario Delmar, Frank Cecchin, Reina B. Tan, Mohamed Chahine, Mohamed Boutjdir

**Author notes:** Corresponding author: Dr. Mohamed Boutjdir, VA New York Harbor Healthcare System, Research and Development Office (151), 800 Poly Place, Brooklyn, NY 11209.

## Abstract

Inherited arrhythmia syndromes are caused by genetic variants that alter cardiac ion channel function. We investigated a complex presentation in a pediatric patient with ventricular tachycardia and conduction abnormalities, harboring a *de novo CACNA1D* (c.3786G>T) variant, and two inherited variants, *the SCN5A* (c.2618C>G), and a *DSP* desmosome (c.1582C>G). The *CACNA1D* variant, which encodes Cav1.3 L-type calcium channel is the focus of this study, because the C-terminus fragment of Cav1.3 has recently been identified as a transcription auto-enhancer of its own gene and able to prevent arrhythmic events in a mouse model of ischemic heart failure. Leveraging this intrinsic property, we hypothesized that the Cav1.3-C-terminus could reverse the arrhythmic events associated with the *CACNA1D* variant.

Patch-clamp and optical mapping experiments demonstrated a loss of Cav1.3 function, characterized by reduced L-type calcium current densities, and decrease of conduction velocity, leading to inducible re-entrant arrhythmias in human induced pluripotent stem cell-cardiomyocytes (hiPSC-CMs). RNA sequencing confirmed this loss-of-function via the downregulation *CACNA1D* gene expression.

Interestingly, Cav1.3-C-terminus treatment of hiPSC-CMs successfully normalized Cav1.3 gene expression, restored calcium currents, and conduction velocity, and prevented the susceptibility to arrhythmias.

These findings highlight the electrophysiological consequences resulting from a *de novo* Cav1.3 variant and demonstrate an important transcriptional role of Cav1.3-C-terminus as a transcriptional regulator and as a promising therapeutic tool to restore normal electrical properties in patients with calcium channels loss of function.

## Introduction

Cardiac arrhythmias represent a critical public health challenge and are a principal driver of sudden cardiac death (SCD), contributing substantially to cardiovascular mortality in the United States. In 2024, cardiovascular disease (CVD) was the underlying cause of almost 1 million deaths^1^. Arrhythmias encompass a heterogeneous set of electrical disorders defined by abnormal impulse initiation or conduction, manifesting as tachyarrhythmias, bradyarrhythmias, or irregular rhythms^2^. These electrophysiological derangements could arise from ion-channel dysfunction due to genetic variants in key regulatory proteins referred to as ion channelopathies ^3^. Sinus node dysfunction is a prototypical form of pacemaker failure, leading to inappropriate bradycardia or pauses that may provoke syncope or SCD^4^. Importantly, ventricular tachyarrhythmias (ventricular tachycardia and ventricular fibrillation) represent the predominant mechanisms underlying SCD^5^. In the cardiovascular system, calcium signaling is crucial for initiation and conduction of the electrical impulses and the subsequent contraction of cardiomyocytes^6,7^. In the adult heart, the calcium signaling is mediated essentially by L-type calcium channels (LTCC), which are composed of Cav1.2, encoded by the *CACNA1C* gene, and Cav1.3 by the *CACNA1D* gene^8^. Cav1.2 is expressed throughout the adult heart, with high expression in the ventricles, while Cav1.3 is found exclusively within the supraventricular tissues, including atria, sinus node (SAN), and atrioventricular node (AVN)^9,10^. Cav1.3 is particularly important in the depolarization during diastole in the SAN and AVN, and in the calcium release from the sarcoplasmic reticulum for the atrial contraction^11^. Disruption of intracellular calcium homeostasis can cause early- and delayed- afterdepolarizations (EADs and DADs, respectively), which act as powerful arrhythmia triggers^12^. Variants in LTCC, mainly gain-of-function, lead to intracellular calcium imbalance, impaired excitation-contraction coupling and systolic dysfunction^13^. Loss of function variants of *CACNA1D* are associated with SAN dysfunction and deafness (SANDD)^14,15^, atrial fibrillation^16^, and bradycardia^14^. In the case of adult human heart failure (HF), Cav1.3 is re-expressed in the ventricles and might serve as a compensatory mechanism to increase contractility^17^.

Previous data show that the C-terminal domain of the Cav1.3 is cleaved and act as an enhancer of its own gene expression^18,19^. Exogenous expression of Cav1.3 was shown to restore normal ejection fraction in mice with HF^20^. In the present proof-of-principle study, we characterized the electrophysiological consequences of a *de novo CACNA1D* variant (c.3786G>T) not previously reported; then leveraged the C-terminus properties as a transcription factor to correct the variant. Our studies were conducted in human induced pluripotent stem cells (hiPSCs) derived from a pediatric patient that presented with intraventricular conduction delay (IVCD), atrioventricular block (AVB), and episodes of ventricular tachycardia.

hiPSCs were guided to differentiate into the three primary types of heart muscle cells: pacemaker-, atrial-, and ventricular-like cardiomyocytes. This approach allowed us to define the contribution of this novel variant to the genesis of chamber-specific arrhythmias. cDNA coding for the Cav1.3 C-terminus was delivered to the hiPSC-CMs via adeno-associated virus serotype 6 (AAV6) viral particles. Previous studies showed that AAV6 is an efficient and a delivery vehicle specific for hiPSC-CMs^21^ that allows for sustained expression of the transgene. AAV6 has been deemed an effective gene delivery system^22,23^ for use in clinical trials^24–26^.

## Methods

### hiPSC culture

Biological samples were handled in accordance with the Declaration of Helsinki with informed consent and standardized approved protocols by a local ethics committee (S14-00862).

Control (CTRL) hiPSC lines were obtained from Stanford University Cardiovascular Institute Biobank. This CTRL hiPSCs is sex- and age-matched to our patient. Patient-specific hiPSCs (referred to as AF13) were derived from peripheral blood mononuclear cells (PBMC) and characterized at the CRCHU of Quebec iPSC Reprogramming Platform^27^.

The hiPSCs were differentiated into pacemaker-, atrial- and ventricular-like cardiomyocytes as previously described^28,29^. Briefly, hiPSCs were plated and cultured on hESC-qualified Matrigel^TM^ (Sigma-Aldrich) at 30,000-50,000 cells/cm^2^ with mTeSR^TM^ for 5 days (StemCells Technologies). The protocol to differentiate the hiPSCs into pacemaker-like cardiomyocytes (hiPSC-PMs) was adapted from Yechikov^28^. Briefly, RPMI Media was supplemented between 0-2 days with 6 μM GSK3 inhibitor (CHIR99021, Tocris), and between 3-5 days with 5 μM Wnt inhibitor (IWR1, Tocris). Small molecule Nodal inhibitor (SB431542, Tocris) was added for 3-5 days. The differentiation process for ventricular and atrial cardiomyocytes was induced with STEMdiff^TM^ Ventricular Cardiomyocyte Differentiation Kit (StemCells Technologies) to produce ventricular-like cardiomyocytes (hiPSC-vCMs) or STEMdiff^TM^ Atrial Cardiomyocyte Differentiation Kit (StemCells Technologies) with all-trans retinoic acid between 2 and 5 days to produce atrial-like cardiomyocytes (hiPSC-aCMs). At the end of the protocol (9 days) the hiPSC-aCMs and hiPSC-vCMs were cultured into STEMdiff^TM^ Cardiomyocyte Maintenance medium (StemCells Technologies) and with RPMI 1640 supplemented with B27 supplement for hiPSC-PMs.

### tsA201 transfection

The human Cav1.3 M1262I variant’s cDNA constructs were developed by site directed mutagenesis. tsA201 cells were cultured in Dulbecco’s Modified Eagle Medium with 10% FBS, 2mM L-glutamine, 100 U/mL of penicillin, and 10 mg/mL of streptomycin (Thermo Fisher Scientific) under standard tissue culture conditions (5% CO2, 37^ο^C). Cells were grown in 35 mm culture plates until 70-80% confluence was reached. Using Lipofectamine 2000 (Invitrogen), as per the manufacturer’s instructions, tsA201 cells were transiently transfected with cDNA plasmids, namely pGFP(+)_hCav1.3 WT or variant (1.2ug), pCDNA3.1_hCACNA2D1 (1.2ug), and pCDNA3.1_hCACNAB2 (1.2ug).

### AAV delivery

AAV6 containing human Cav1.3-C-terminus (AA N1840-L2137) was generated by the Neurophotonics, CERVO Brain Research Centre Molecular Tools Platform. The hiPSC-CMs were infected with 5.10^3^ v.g. per cell. After 48 hours, the virus was removed, and the experiments were performed after 4 or 5 days.

### Gene Expression Analysis

#### Quantitative reverse transcription polymerase chain reaction

For RNA extraction, a 30-day-old beating hiPSC-CMs monolayer was used. Total RNA was isolated using a NucleoSpin RNA kit (Macherey-Nagel) followed by a reverse transcription to obtain cDNA using the Transcriptor Universal cDNA Master Mix (Sigma-Aldrich), according to manufacturer’s’ protocols. For detection of the gene of interest, cDNA was amplified by the PowerUp SYBR Green Master Mix (Applied Biosystems) for qRT-PCR. cDNA amplifications were performed with the Quant Studio 5 from Applied Biosystems. Human Large Ribosomal Protein (RPLPo) was used as a reference gene. The genes of interest studied by RT-qPCR included *CACNA1C*, *CACNA1D*, *SCN5A*, *HCN4*, and *DSP*.

#### RNA-seq

The RNA-seq experiments were performed by AZENTA Life sciences. Briefly, Total RNA was extracted from hiPSC-CMs frozen cell pellets using Qiagen RNeasy Plus Universal Mini kit following manufacturer’s instructions (Qiagen). RNA samples were quantified using Qubit 2.0 Fluorometer (Life Technologies), and RNA integrity was checked using Agilent TapeStation 4200 (Agilent Technologies). RNA sequencing libraries were prepared using the NEBNext Ultra II RNA Library Prep Kit for Illumina and the NEBNext Poly(A) mRNA Magnetic Isolation Module following the manufacturer’s instructions (NEB). Briefly, mRNAs were initially enriched with Oligo(T) beads. Enriched mRNAs were fragmented for 15 minutes at 94°C. First-strand and second-strand cDNA were subsequently synthesized. cDNA fragments were end-repaired and adenylated at 3’ ends, and universal adapters were ligated to cDNA fragments, followed by index addition and library enrichment by PCR with limited cycles. The sequencing library was validated on the Agilent TapeStation (Agilent Technologies) and quantified by using a Qubit 2.0 Fluorometer (Invitrogen) as well as by quantitative PCR (KAPA Biosystems). The sequencing libraries were clustered on a flowcell. After clustering, the flowcell was loaded on the Illumina NovaSeq X Plus instrument according to the manufacturer’s instructions. The samples were sequenced using a 2x150bp Paired-End (PE) configuration, targeting approximately ∼30M reads/sample. The control software conducted image analysis and base calling. Sequence reads were trimmed to remove possible adapter sequences and nucleotides of poor quality using Trimmomatic v.0.36. The trimmed reads were mapped to the *Homo sapiens GRCh38* reference genome available on ENSEMBL using the STAR aligner v.2.5.2b. STAR is a splice aligner that detects splice junctions and incorporates them to help align entire read sequences. Mapping .bam files were used to generate hit counts for genes and exons.

Unique gene hit counts were calculated by using featureCounts from the Subread package v.1.5.2. The hit counts were summarized and reported using the gene_id feature in the annotation file. Only unique reads that fell within exon regions were counted. Genes with an adjusted p-value < 0.05 and absolute log2 fold change > 1 were called as differentially expressed genes.

### Immunofluorescence

hiPSC-CMs were dissociated at D23 using TrypLE^TM^ and were seeded on Matrigel-coated 35-mm µ-dish glass bottom (Ibidi) at a density of 30,000 cells/cm^2^. At D30, the cells were fixed with 4% paraformaldehyde and permeabilized in a PBS solution containing 0.1% Triton X-100 for 15 min. The cells were saturated using Normal Goat Blocking Buffer (Elabscience) for 30 min. hiPSC-CMs were stained overnight at 4°C using PBS containing the following primary antibodies: mouse anti-alpha actinin (1:200, Ca# A7732, Sigma) and rabbit anti-Cav1.3 (1:200, Ca# ACC-005, Alomone). The secondary antibodies, goat anti-mouse IgG Elab Fluor 488 (1:200, Cat# E-AB-1056, Elabscience) and goat anti-rabbit IgG Elab Fluor 594 (1:200, Cat# E-AB-1060, Elabscience) were then incubated for 60 min at room temperature in the dark. The nucleus was labelled using DAPI Reagent (Cat# E-IR-R103, Elabscience) for 5 min. Cells were observed at 63x objective using a Zeiss LSM800 confocal microscope (Zeiss) and examined with ZEN software.

### hiPSC-CMs contraction analysis

The hiPSC-CMs contractile function was performed using patented, custom-made video analysis software, as previously published^30,31^. The hiPSC-CMs monolayer (without dissociation) in twelve-well plates were placed on the stage of an inverted microscope equipped with a 20x objective. Phase contrast videos were acquired at 60 frames per second. Videos were then processed: TIFF images were extracted, and contrasted particle displacement was tracked frame by frame for each video. The displacement of each contrasted particle was then processed through the time duration, resulting in a curve of the displacement as a function of time. Areas with similar contractile behavior were clustered and contractile parameters quantified by a Matlab script.

### Electrophysiology

#### Optical mapping

hiPSC-CMs were enzymatically dissociated (TrypLE^TM^) and seeded onto hESC-qualified Matrigel-coated 12 mm TC coverslips in a 24-well plate between 12 and 15 days after the differentiation. At 30-35 days, the hiPSC-CMs monolayer were stained with Rhod-2 AM (5 μM, Abcam), a calcium indicator, and incubated at 37°C for 30 min. The imaging solution contained (in mmol/L): 154 NaCl, 5.6 KCl, 2 CaCl2, 1 MgCl2, 8 glucose and 10 HEPES; pH was adjusted at 7.3 with KOH. An epifluorescence macroscope, equipped with CMOS N256 cameras (MiCAM03, Brainvision), was employed for the optical mapping system to capture calcium transients at a rate of 500 frames per second. Calcium transient was used to maximize signal to noise ration compared to voltage. The hiPSC-CMs monolayer were maintained at 37°C using a controlled heating plate (MappingLab) and were electrically stimulated using two Teflon-coated silver electrodes positioned at the lower edge of the monolayer. A 385-stimulus generator (WPI) was used to deliver 80 mA current injections. S1 stimuli were delivered to the monolayers at 1.0 and 2.0 Hz. Premature stimuli (S2-S4) were delivered to mimic ectopic beats, as previously described^32^. The raw optical signals were processed and analyzed using Brainvision Workbench software (Brainvision, SciMedia).

#### Patch Clamp

Patch clamp experiments were performed at room temperature using an Axopatch 200B amplifier (Axon Instruments). Pipettes were made from borosilicate glass capillaries, and fire polished.

Action potentials (APs) were evaluated in current clamp mode using the whole cell configuration. APs were elicited at 1 Hz following injection of a 3 ms, 20-1500 pA rectangular current pulse. The patch pipettes (resistance 2-3 MOhms) were filled with a solution containing (in mmol/L): 10 NaCl, 122 KCl, 1 MgCl2, 1 EGTA and 10 HEPES; pH was adjusted at 7.3 with KOH. The bath solution (external current clamp) was composed of (in mmol/L): 154 NaCl, 5.6 KCl, 2 CaCl2, 1 MgCl2, 8 glucose and 10 HEPES; the pH was adjusted at 7.3 with NaOH.

For calcium currents (ICaL), the patch pipettes (resistance 1-2 MOhms) were filled with (in mmol/L): 25 NaCl, 105 CsCl, 1 MgCl2, 10 EGTA and 10 HEPES; pH adjusted at 7.3 with CsOH. The bath solution consisted of (in mmol/L): 100 NaCl, 5 CsCl, 5 CaCl2, 40 NMDG, 1 MgCl2, 10 glucose, 10 HEPES and 15 TEA-Cl; pH was adjusted at 7.3 with methanethiosulfonic acid. Calcium currents were obtained using 200 ms square steps to +35mV from a holding potential of -40 mV at 5 s intervals.

For sodium currents (INa), patch electrodes (resistance 1-2 MOhm) were filled (in mmol/L): 105 CsF, 35 NaCl, 10 EGTA, 10 HEPES and pH adjusted to 7.2 with CsOH. The bath solution consisted of (in mmol/L): 105 NMDG, 35 NaCl, 2 KCl, 10 HEPES, 1 MgCl2, 1.5 CaCl2, 10 TEA-Cl and 0.5 Nimodipine with pH 7.4 adjusted with methanethiosulfonic acid. INa were obtained using 50 ms pulses from -90 mV to +45mV in +5mV increments.

### Data analysis and statistics

Optical mapping and patch clamp results were analyzed using Brainvision software and Clampfit (pCLAMP v10; Molecular devices), respectively. The contractile function was analyzed using custom-written Matlab programs. The code is available upon request. A normality test (Agostino and Pearson omnibus normality test) was always used to determine whether data followed a normal distribution. Data processing and statistical analyses were conducted with Prism 10. Mean with s.e.m. was calculated from experiments obtained from at least three independent differentiations. One- or two-way ANOVA test was used to assess significance (ns p > 0.05; *p < 0.05, **p < 0.01, ***p < 0.001).

## Results

### Patient description

The hiPSCs cell line was derived from PBMCs of a 13-year-old female patient (AF13). The patient initially presented at 7 years old with symptomatic ventricular tachycardia (VT) **Table 1** and **Figure 1**. The patient inherited two variants, *SCN5A* (c.2618C>G; p.Ser873*, inducing a stop codon) and *DSP* (c.1582C>G; p.Gln528Glu) out of three variants from her mother. The patient also harbored a new variant on the *CACNA1D* (c.3786G>T; p.Met1262Ile) gene. This variant results in amino acid substitution M1262I, topologically located in the second transmembrane segment of Domain IV (**Figure 1A, B and C**). There was no evidence of Brugada pattern on her ECGs. Holter monitor showed sinus bradycardia and frequent runs of VT. Echocardiogram showed a structurally normal heart with normal biventricular function. Cardiac MRI showed mild right ventricular dilation with normal function. She failed sotalol therapy and flecainide resulted in significant QRS prolongation. She was managed on amiodarone with some improvement however patient would self-discontinue medication. Trial of quinidine resulted in significant improvement of VT burden; however, patient was non complaint with medications. She underwent an electrophysiology study that demonstrated bundle branch reentrant VT which was successfully eliminated with radiofrequency catheter ablation. There was recurrence of VT 3-months post ablation albeit with at least 90% decrease in VT burden.

**Figure 1:**
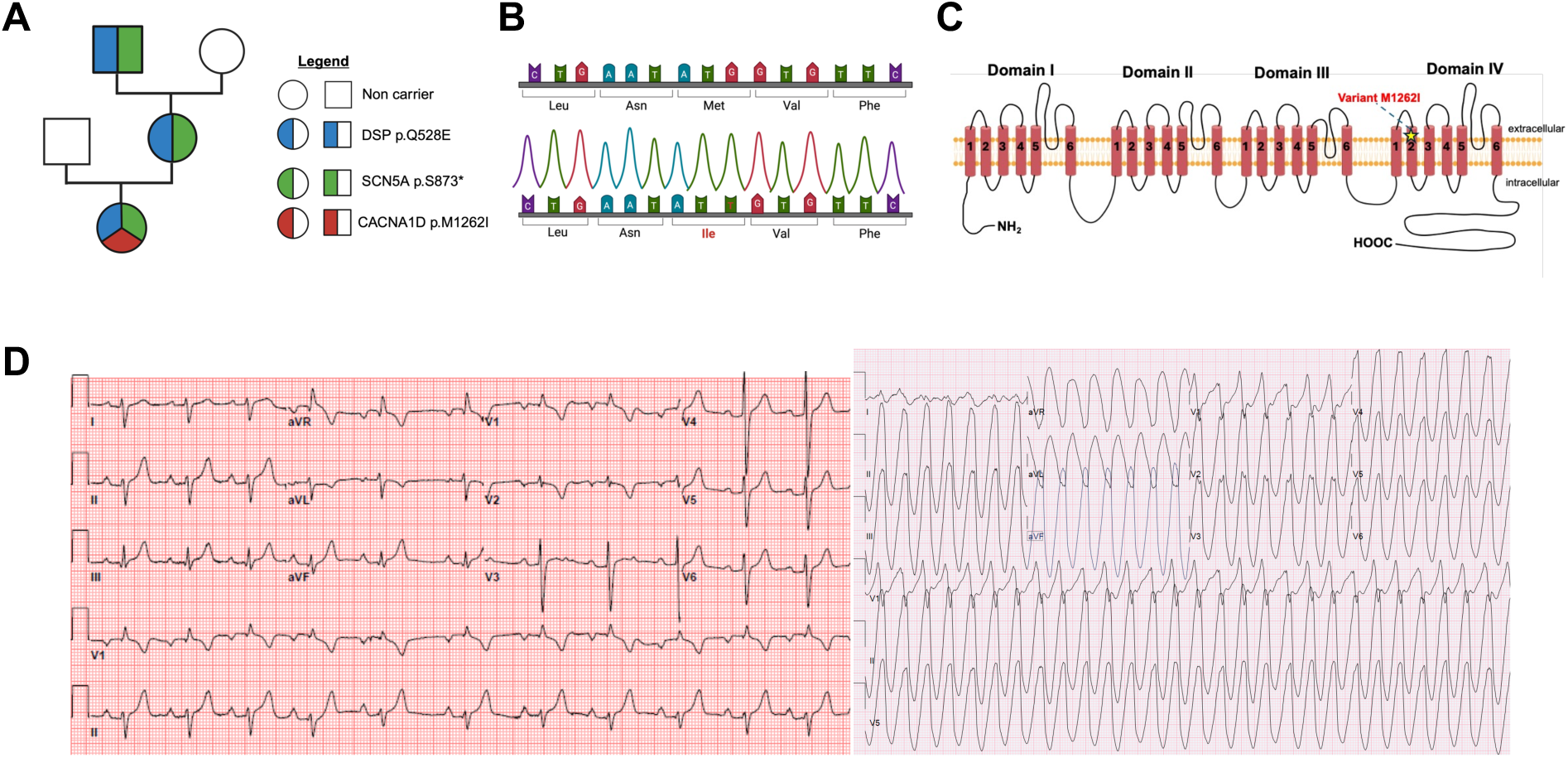
Clinical presentation. **(A)** Family pedigree indicating the carried variants. **(B)** Schematic chromatogram of the patient’s DNA showing the *CACNA1D* variant responsible for the M1262I substitution. **(C)** Schematic representation of Cav1.3 protein structure with the localization of the new variant in transmembrane segment 2 of Domain IV. **(D)** Twelve-lead surface ECG illustrating the patient’s electrical profile.

**Table 1:** Clinical presentation of the patient and her mother

|  | Genetics | Age | Rhythm | ECG | Morphology | Arrhythmia | Medication |
| --- | --- | --- | --- | --- | --- | --- | --- |
| Daughter | SCN5A<br>p.S873*<br>DSP<br>p.Q528E<br>CACNA1D<br>p.M1262I | 13 yrs | Sinus<br>Bradycardia<br>Minimum:<br>47bpm<br>Maximum:<br>205bpm<br>Average:<br>80bpm | Wide QRS<br>Prolong First<br>degree AV<br>block | Mild RV<br>volume<br>increase<br>Normal<br>function<br>No HF sign | Episode of<br>Ventricular<br>tachycardia | Amiodarone |
| Mother | SCN5A<br>p.S873*<br>DSP<br>p.Q528E | 38 yrs | Normal<br>sinus rhythm | Diagnosed<br>type 1 Brugada<br>(asymptomatic) | N.A. | No<br>arrythmia | No<br>medication |

**Table 2:** Action Potential parameters recording in hiPSC-CMs. Value are means±SEM. CTRL: Control hiPSC-CMs; AF13: hiPSC-CMs carrying Cav1.3 variant; +AAV: AF13 + AAV6 Cav1.3-C-terminus treatment. * CTRL versus AF13, # AF13 versus +Cterm, § CTRL versus +Cterm; *p < 0.05, **, ##, §§p < 0.01, ***, ###p < 0.001 (Kruskal-Wallis Test multiple comparisons test).

|  |  | APD<sub>90</sub> (ms) | APD<sub>50</sub> (ms) | APD<sub>30</sub> (ms) | Amplitude (mV) | dV/dt<sub>max</sub> (mV/ms) |
| --- | --- | --- | --- | --- | --- | --- |
| Pacemaker-like | CTRL (n=18) | 114.8±10.3 | 47.6±5.9 | 29.4±4.7 | 99.0±2.7 | 32.1±4.4 |
|  | AF13 (n=17) | 95.7±11.1 | 36.0±4.7 | 13.4±2.3 *** | 52.1±3.6 *** | 9.0±1.0 *** |
|  | +Cterm (n=18) | 108.3±8.8 | 38.0±5.5 | 19.4±4.6 | 89.9±5.2 ## | 21.67±2.6 ## |
| Atrial-like | CTRL (n=27) | 272.3±16.5 | 150.3±10.8 | 95.1±9.7 | 123.2±3.9 | 72.6±7.9 |
|  | AF13 (n=26) | 190.2±11.7 ** | 97.9±9.1 ** | 70.0±7.5 | 100.2±3.1 *** | 24.2±4.1 ** |
|  | +Cterm (n=28) | 278.9±6.9 ### | 183.9±7.9 ### | 122.2±6.6 ### § | 129.2±3.2 ### | 89.7±7.2 ### |
| Ventricular-like | CTRL (n=30) | 435.0±19.1 | 300.4±18.2 | 225.9±13.8 | 110.6±2.9 | 36.4±5.0 |
|  | AF13 (n=22) | 312.1±12.8 *** | 188.7±7.9 *** | 148.2±7.3 *** | 97.8±2.3 * | 22.7±3.3 * |
|  | +Cterm (n=22) | 452.4±21.2 ### | 338.9±14.5 ### | 259.0±13.1 ### | 128.1±1.9 ### §§ | 77.4±4.3 ### §§ |

**Table 3:** Contractile parameters of hiPSC-CMs Value are means±SEM. CTRL: Control hiPSC-CMs; AF13: hiPSC-CMs carrying Cav1.3 variant; +AAV: AF13 + AAV6 Cav1.3-C-terminus treatment. #p < 0.05, **, ##p < 0.01, ***, ###p < 0.001, ****, ####p < 0.0001. (Kruskal-Wallis Test multiple comparisons test).

|  |  | Beating rate (bpm) | Contraction duration (ms) | Relaxation duration (ms) | Homogeneity |
| --- | --- | --- | --- | --- | --- |
| Pacemaker-like | CTRL (n=43) | 78.9±2.9 | 186.5±9.0 | 510.5±17.2 | 1.1±0.05 |
|  | AF13 (n=41) | 30.9±1.4 **** | 128.9±6.3 *** | 533.2±31.7 | 0.4±0.06 **** |
|  | +Cterm (n=38) | 83.3±3.8 #### | 190.9±7.0 ### | 536.9±17.9 | 1.2±0.08 #### |
| Atrial-like | CTRL (n=42) | 79.8±4.2 | 262.4±13.2 | 655.5±30.8 | 1.1±0.1 |
|  | AF13 (n=36) | 43.3±3.9 ** | 129.1±5.2 ** | 445.0±19.6 ** | 0.2±0.02 ** |
|  | +Cterm (n=35) | 70.7±2.9 # | 279.6±17.7 ## | 688.6±17.2 ## | 0.9±0.07 ## |
| Ventricular-like | CTRL (n=32) | 62.1±4.0 | 339.2±26.1 | 622.3±24.4 | 1.4±0.07 |
|  | AF13 (n=43) | 26.1±1.1 *** | 226.8±9.4 ** | 559.5±20.3 | 0.34±0.07 **** |
|  | +Cterm (n=36) | 56.9±3.3 ### | 343.7±14.1 #### | 682.1±19.0 ### | 0.86±0.06 #### |

**Table 4:** Optical mapping data from hiPSC-CMs monolayer

|  |  | Conduction Velocity<br>(cm/s) | Upstroke Velocity<br>(Unit/ms) | Ca<sup>2+</sup> Decay Time (ms) |
| --- | --- | --- | --- | --- |
| Pacemaker-like | CTRL<br>(n=5) | 4.6±0.5 | 52±3.9 | 242.8±14.9 |
|  | AF13<br>(n=5) | 2.2±0.2 * | 22.4±3.7 * | 155±10.8 * |
|  | +Cterm<br>(n=5) | 4.8±0.4 # | 52.4±7.1 # | 237±23 # |
| Atrial-like | CTRL<br>(n=5) | 6.4±0.8 | 99.2±10 | 463.4±28.3 |
|  | AF13<br>(n=5) | 3.4±0.3 * | 50.4±6.9 * | 237.6±15.2 * |
|  | +Cterm<br>(n=5) | 6.9±0.9 # | 104.6±9.5 # | 487.6±47.6 # |
| Ventricular-like | CTRL<br>(n=5) | 16.1±0.6 | 79.4±4.6 | 709.6±21.4 |
|  | AF13<br>(n=5) | 7.1±1.3 ** | 30.1±7.9 ** | 447.0±48.7 * |
|  | +Cterm<br>(n=5) | 14.9±0.6 | 69.4±1.7 | 734.0±16 ## |

### Molecular phenotypic hallmarks observed in Cav1.3 hiPSC-CMs

To investigate the expression and localization of the Cav1.3 channel, we performed immunofluorescence on hiPSCs from the patient (AF13) and control (CTRL) that were successfully differentiated into three cardiomyocyte subtypes. Immunostaining for Cav1.3 revealed a perinuclear and membrane localization in CTRL hiPSC-aCMs (atrial), whereas no expression was observed in the AF13 cells (**Figure 2A and B**, respectively). Notably, the introduction of an AAV6 Cav1.3-C-terminus resulted in the re-expression of Cav1.3 in AF13 hiPSC-aCMs, with a localization pattern like that of the CTRL group (**Figure 2C**). Several cardiomyocyte specific genes (*CACNA1D, CACNA1C, SCN5A,* and *DSP*) were also evaluated using RT-qPCR. In hiPSC-aCMs, a significant 3-fold decrease in the expression of the *CACNA1D* gene was observed in AF13 compared to CTRL. The AAV6 Cav1.3-C-terminus treatment successfully recovered this expression to control levels (**Figure 2D**). Conversely, expression of the *CACNA1C* was present in all three groups of hiPSC-aCMs with no significant differences observed (**Figure 2E**). Furthermore, the expression levels of the sodium channel gene, *SCN5A,* tended to be reduced (30% in AF13 and +Cterm group compared to CTRL) without statistical significance, and desmoplakin, *DSP*, remained unchanged across all three groups (**Figure 2F and G**). Interestingly, the same Cav1.3 localization and expression pattern were found in hiPSC-PMs (pacemaker) and hiPSC-vCMs (ventricular) **Supplemental Figure 1 and 2**). The gene expression of *HCN4* coding for the pacemaker current, If, in hiPSC-PMs, was not altered for each condition (**Supplemental Figure 1E**). The expression of the *DSP* gene, which is associated with cell adhesion, and *SCN5A* have also a tendency to be reduced in AF13 hiPSC-vCMs compared to CTRL (**Supplemental Figure 2F and G**). It is also important to note that the cardiomyocyte identity of the cells was validated through α-actinin staining, where the hiPSC-PMs exhibited a more elongated morphology compared to the other two subtypes, which is consistent with a pacemaker cell phenotype^34^.

**Figure 2:**
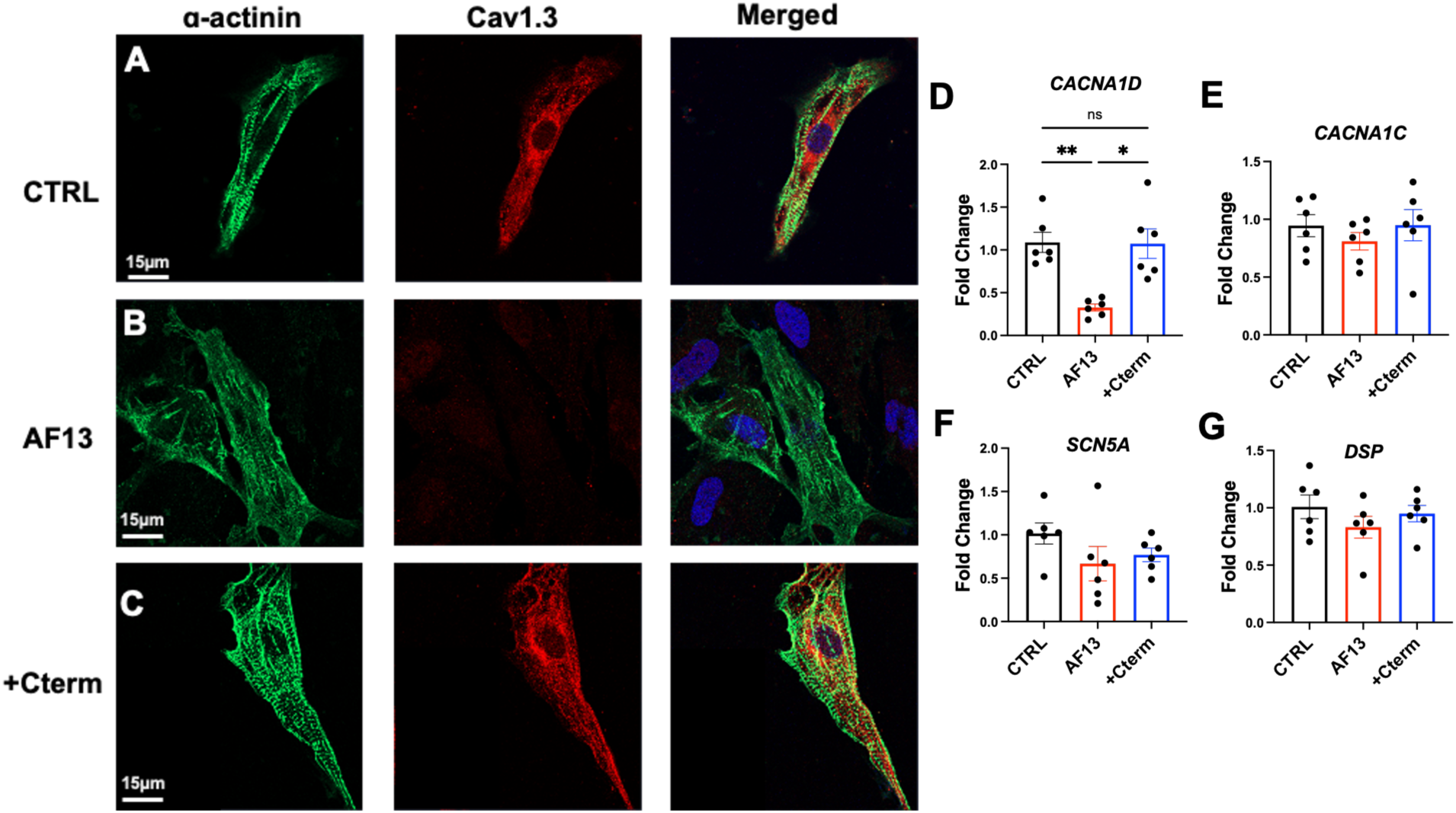
Mislocalization of Cav1.3 and decrease of *CACNA1D* expression, rescued by AAV6 Cav1.3-C-terminus treatment in atrial-like hiPSC-aCMs. Representative images of atrial-like hiPSC-CMs (hiPSC-aCMs) stained with α-actinin (green), Cav1.3 (red) and nuclei (DAPI, cyan) in CTRL (**A**), AF13 (**B**) and after treatment with AAV6 Cav1.3-C-terminus (**C**). Immunofluorescence images were acquired using Zeiss LSM800 confocal microscope and processed with ZEN software (Zeiss). RT-qPCR analysis of several genes implicated in cellular excitability (*CACNA1D*, *CACNA1C,* and *SCN5A* (**D-F**)), cardiomyocytes adhesion (*DSP* (G)). N = 6 for gene expression analysis. Results are shown with the mean±s.e.m. * p<0.05, ** p<0.01, ns not statically significant (Kruskal-Wallis test). CTRL: Control hiPSC-aCMs; AF13: hiPSC-aCMs carrying Cav1.3 variant; +Cterm: AF13 + AAV6 Cav1.3-C-terminus treatment

Next, we performed RNA-seq for a more detailed, unbiased analysis of the expression levels of *CACNA1D* and other genes both prior to and following AAV6 Cav1.3-C-terminus treatment.

In hiPSC-aCMs, the heat map and volcano plot data revealed that 266 genes were downregulated and 284 were upregulated in the AF13 compared to CTRL (**Figure 3A and B**). The upregulated genes included those associated with potassium ion transport (*KCNJ13*, *KCND3*, and *KCNK1*), cardiac muscle hypertrophy (*LMCD1*), and apoptotic signaling (*MMP9*). In contrast, the downregulated genes were found to be involved in calcium ion transport and the cardiac conduction system (*CACNA1D* and *HCN1*), along with the regulation of heart contraction (*MYOM2* and *MYH6*) (**Figure 3C and D**). These results substantiate the downregulation of *CACNA1D* expression. Following the treatment of AF13 hiPSC-aCMs with AAV6 Cav1.3-C-terminus, the heat map and volcano plot data revealed that 1,381 genes were downregulated and 839 were upregulated (**Figure 3E and F**). The upregulated genes included calcium ion transport (*CACNA1D*), and those involved in the regulation of gene expression (*TAPBP* and *TC2N*), heart contraction (*MYL2*, *TNNI3*, and *PLN*), sarcomere organization (*CASQ2* and *MYPN*). The downregulated genes were associated with potassium ion transport (*KNCQ2*, *KCNJ13*, *KCNE4*, and *KCNK2*) and sodium ion transport (*SLC13A4*, *SCN7A*, and *SCN3A*) (**Figure 3G and H**). These RNA-seq findings confirm the upregulation of the *CACNA1D* gene upon AAV6 Cav1.3-C-terminus treatment and indicate enhanced cardiomyocyte sarcomere formation. Comparable results were observed in both pacemaker and ventricular conditions (**Supplemental Figure 3**).

**Figure 3:**
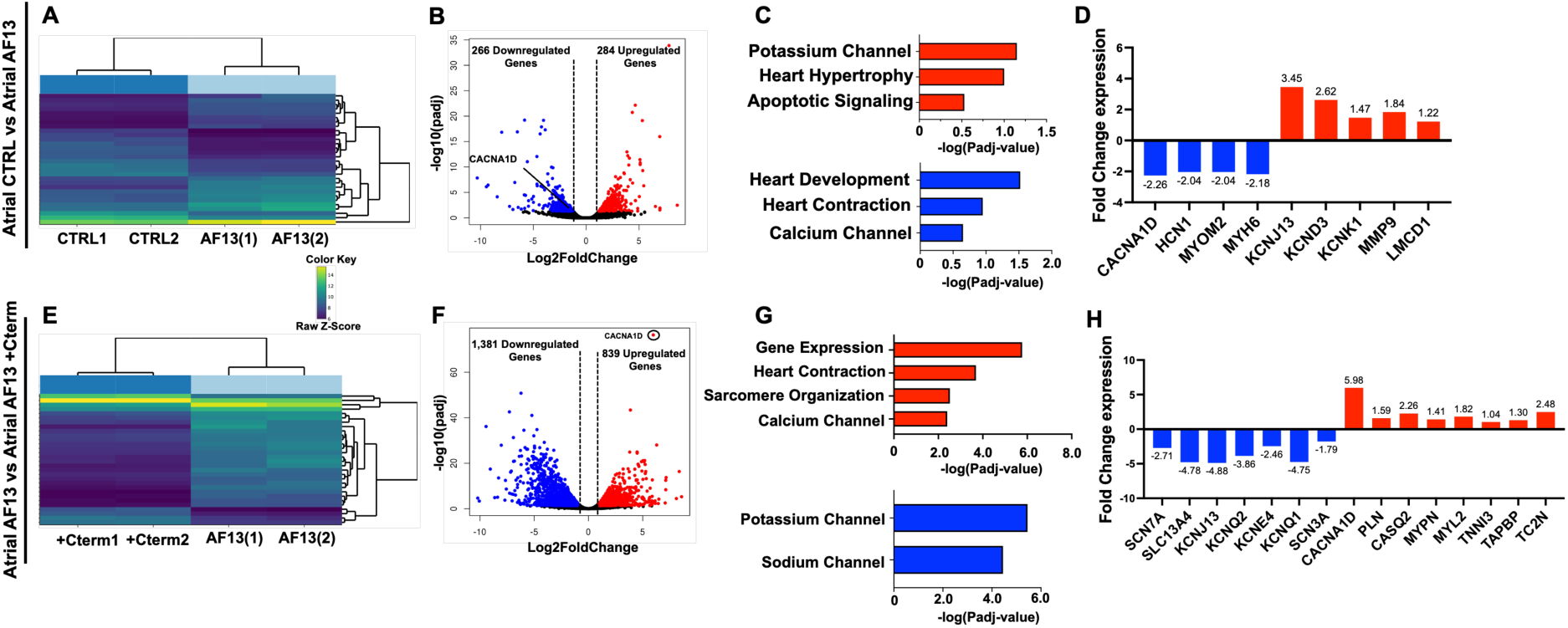
Transcriptomic analysis of human induced pluripotent stem cell atrial-like cardiomyocytes (hiPSC-aCMs). **(A)** Heat map plot of the top 30 DEGs between CTRL and AF13 condition. **(B)** Volcano plot of downregulated (blue) and upregulated (red) genes in CTRL and AF13. **(C)** Gene ontology biological analysis of DEGs in CTRL and AF13. **(D)** Fold change in expression of key genes. **(E)** Heat map plot of the top 30 DEGs between AF13 and after AAV6 Cav1.3-C-terminus incubation (+Cterm). **(F)** Volcano plot of downregulated (blue) and upregulated (red) genes in AF13 and +Cterm **(G)** Gene ontology biological process analysis of DEGs in AF13 and +Cterm. **(H)** Fold change in expression of key genes. The Wald test was used to generate p-values and log2 fold changes. Genes with an adjusted p-value < 0.05 and absolute log2 fold change > 1 were called as differentially expressed genes. CTRL: Control hiPSC-aCMs; AF13: hiPSC-aCMs carrying Cav1.3 variant; +Cterm: AF13 + AAV6 Cav1.3-C-terminus treatment.

### Effect of Cav1.3 variant and AAV6 Cav1.3-C-terminus treatment on ionic current properties

The properties of APs were studied in pacemaker-, atrial- and ventricular-like hiPSC-CMs by the patch-clamp technique. Several AP parameters were analyzed, including AP duration at 90% of repolarization (APD90) and 50% of repolarization (APD50), amplitude and the maximal upstroke velocity (dV/dtmax). These AP parameters were affected by the variant but corrected by the AAV6 Cav1.3-C-terminus treatment (**Figure 4A, B and C**). In hiPSC-PMs, the APD90 and APD50 were not significantly changed between all conditions (**Figure 4D and E**). However, in both hiPSC-aCMs and hiPSC-vCMs the APD90 was shortened by 30% and 28%, respectively, in AF13 compared to CTRL. Interestingly, the AAV6 Cav1.3-C-terminus treatment normalized the APD90 (**Figure 4D**). The same results were observed for APD50 (**Figure 4E**). The AP amplitude was also decreased by 47% in hiPSC-PMs, 18% in hiPSC-aCMs and 12% in hiPSC-vCMs in AF13 compared to CTRL (**Figure 4F**). The AAV6 Cav1.3-C-terminus normalized the AP amplitude in hiPSC-PMs and hiPSC-aCMs, but it increased the AP amplitude in hiPSC-vCMs (**Figure 4F**). The dV/dtmax was decreased in all cardiomyocyte’s subtypes between CTRL and AF13. Like the previous AP parameters, the AAV6 Cav1.3-C-terminus corrected the maximal upstroke velocity for each cardiomyocyte subtype (**Figure 4G**).

**Figure 4:**
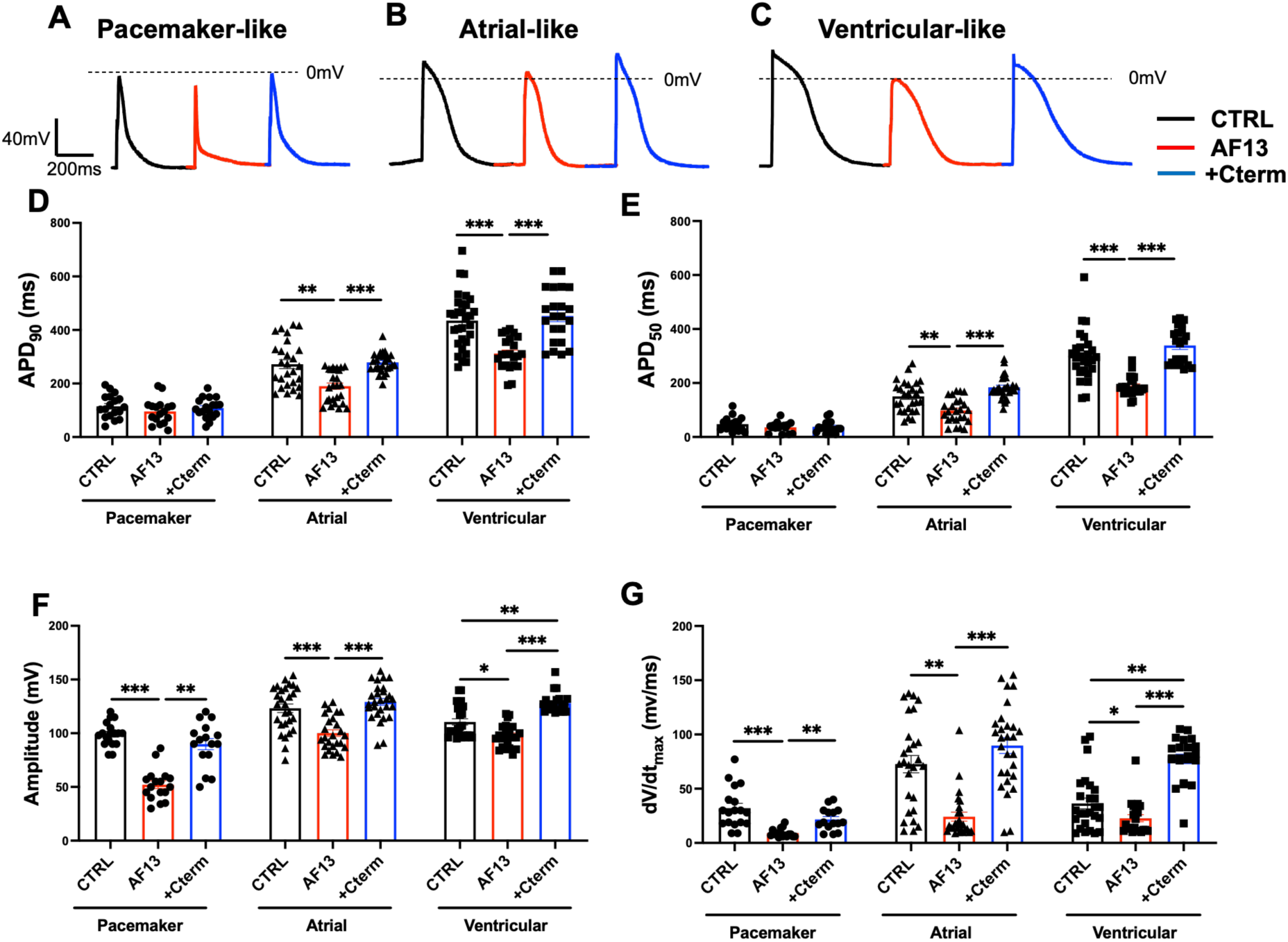
Characterization of action potential in cardiomyocytes subtypes by single cell patch-clamp. **(A)** Raw traces of APs in CTRL (black; n=18), AF13 (red; n =17) and after AAV6 Cav1.3-C-terminus treatment (blue; n=18) in hiPSC-pacemaker-like cardiomyocytes (hiPSC-PMs). **(B)** Raw traces of APs in CTRL (black, n=27), AF13 (red; n=26) and after AAV6 Cav1.3-C-terminus treatment (blue; n=28) in hiPSC-atrial-like cardiomyocytes (hiPSC-aCMs). **(C)** Raw traces of APs in CTRL (black; n=30), AF13 (red; n=22) and after AAV6 Cav1.3-C-terminus treatment (blue; n=22) in hiPSC-ventricular-like cardiomyocytes (hiPSC-aCMs). **(D)** Measurements of AP duration at 90% of repolarization (APD90) for all cardiomyocytes subtypes and conditions. **(E)** Measurements of AP duration at 50% of repolarization (APD50) for all cardiomyocytes subtypes and conditions. **(F)** Values of AP amplitude and **(G)** values of maximal AP upstroke (dV/dtmax) for all cardiomyocytes subtypes and condition. N = 3 independent differentiations. Error bars, s.e.m. Kruskal-Wallis Test. *p < 0.05, **p < 0.01, ***p < 0.001. CTRL: Control hiPSC-CMs; AF13: hiPSC-CMs carrying Cav1.3 variant; +Cterm: AF13 + AAV6 Cav1.3-C-terminus treatment.

We further explored the effects of the *de novo CACNA1D* variant on the biophysical properties of L-type calcium currents, ICaL, by voltage-clamp mode. Firstly, the AF13 hiPSC-PMs showed a lower ICaL densities than the CTRL, consistent with the downregulation of *CACNA1D* gene by RT-PCR and RNA-seq and that the AAV6 Cav1.3-C-terminus treatment normalizes this current (**Figure 5A, B and C**). The ICaL current density-voltage (I/V) relationship revealed a significant 3.0-fold decrease, while AAV6 Cav1.3-C-terminus treatment revealed a rescue comparable to CTRL without activation and inactivation kinetics modification (**Figure 5D and E**). The hiPSC-aCMs also exhibited a decreased of ICaL between CTRL and AF13, which was also restored by the AAV6 Cav1.3-C-terminus treatment (**Figure 5F, G and H**). The I/V relationship revealed a 2.0-fold decrease, while AAV6 Cav1.3-C-terminus treatment normalized the ICaL compared to CTRL, without kinetics modification (**Figure 5I and J**). Similarly, in hiPSC-vCMs, ICaL was reduced between CTRL and AF13 and normalized by the AAV6 Cav1.3-C-terminus treatment (**Figure 5K, L and M**). The I/V relationship revealed a significant 1.6-fold decrease, while AAV6 Cav1.3-C-terminus treatment corrected the loss of function compared to CTRL, with an inactivation kinetics slightly modified, but not statistically significant when compared to AAV6 Cav1.3-C-terminus (**Figure 5N and O**). We also confirmed the loss of function in Cav1.3 due to this variant in tsA201 cells by transfection (**Supplemental Figure 4**). It is important to note that we confirmed and validated the expression of Cav1.3 in hiPSC-vCMs by applying a specific inhibitor of Cav1.2, calciseptine^35^ (**Supplemental Figure 5**). The application of calciseptine (1µM) reduced the ICaL by 60%, leaving 40% of ICaL carried by the Cav1.3 (**Supplemental Figure 5**). Collectively, these results demonstrated a loss of function in ICaL in the hiPSC-CMs carrying the variant and the AAV6 Cav1.3-C-terminus treatment remarkably restoring ICaL to normal values in these cells.

**Figure 5:**
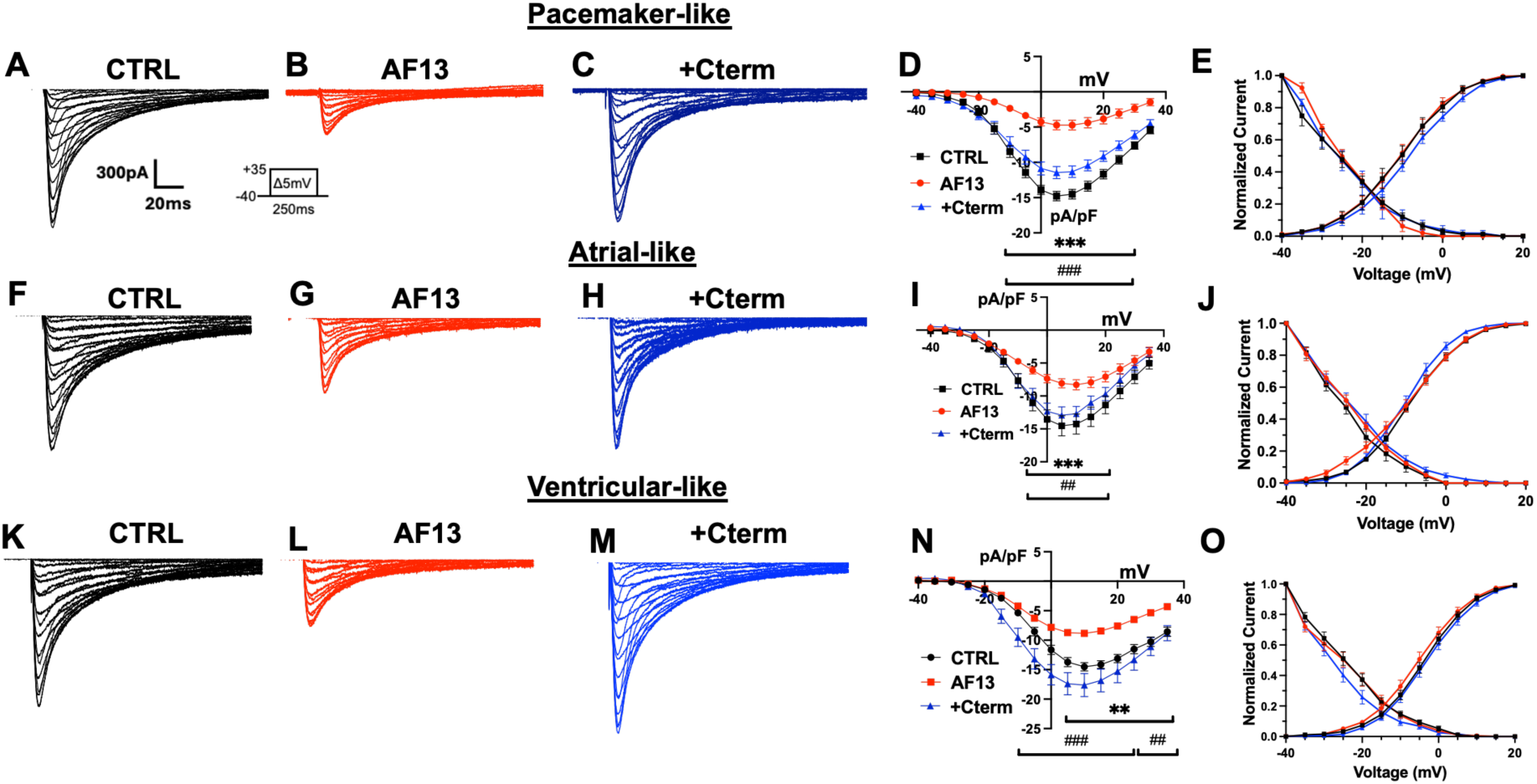
Biophysical properties of L-type calcium channels in hiPSC pacemaker-, atrial- and ventricular-like subtypes. Representative calcium current recorded in (**A**) CTRL (black; n=19); (**B**) AF13 (red; n=15) and (**C**) after AAV6 Cav1.3-C-terminus treatment (+Cterm) (blue; n=17) in hiPSC-PMs. (**D**) Normalized Current/Voltage relationship curves, and (**E**) steady-state activation and inactivation in hiPSC-PMs. Representative calcium current recorded in (**F**) CTRL (black; n=17); (G) AF13 (red; n=18) and (**H**) +Cterm (blue; n=16) in hiPSC-aCMs. (I) Normalized Current/Voltage curves, and (**J**) steady-state activation and inactivation in hiPSC-aCMs. Representative calcium current recorded in (**K**) CTRL (black; n=18); (L) AF13 (red; n=15) and (**M**) +Cterm (blue; n=15) in hiPSC-vCMs. (N) Normalized Current/Voltage curves, and (O) steady-state activation and inactivation in hiPSC-vCMs. N = 3 independent differentiations. Error bars, s.e.m. A two-way ANOVA with Sidak multiple comparisons test. **p < 0.01, ***p < 0.001 (CTRL vs. AF13), and ##p < 0.01, ##p < 0.001. CTRL: Control hiPSC-CMs; AF13: hiPSC-CMs carrying Cav1.3 variant; +Cterm: AF13 + AAV6 Cav1.3-C-terminus treatment.

We also performed INa recording in hiPSC-aCMs and hiPSC-vCMS to elucidate the pathogenicity of the *SCN5A* variant with a stop codon. No significance was found in INa densities between CTRL and AF13 in hiPSC-aCMs subtypes (**Supplemental Figure 6 A, B and C**). Interestingly, the INa density was significantly decrease by 2-fold in AF13 compared to CTRL in hiPSC-vCMs (**Supplemental Figure 6D, E and F).**

### Alterations of monolayer hiPSC-CMs contractile and propagation properties by the Cav1.3 variant and rescue by Cav1.3 C-terminus treatment

The impact of the *CACNA1D* variant on the contractile function of hiPSC-CMs monolayer from CTRL, AF13, and AAV6 Cav1.3-C-terminus treated groups were assessed using video analysis (**Figure 6**). Examples of the contraction profiles are shown in **Figure 6A-C**, with key parameters such as spontaneous beating rate, contraction and relaxation duration, and homogeneity representing the uniform contraction of the monolayer. The spontaneous beating rate and contraction duration were lower in the AF13 patient’s cells compared to CTRL cells across all cardiomyocyte subtypes. Interestingly, AAV6 Cav1.3-C-terminus treatment normalized both the spontaneous beating rate and the contraction duration in every cardiomyocyte subtype (**Figure 6D and E**). However, the relaxation duration remained unchanged in hiPSC-PMs and hiPSC-vCMs (**Figure 6F**). Only in hiPSC-aCMs, did we observe a decrease in relaxation duration in the AF13 cells, which was subsequently normalized following AAV6 Cav1.3-C-terminus treatment (**Figure 6F**). The homogeneity was lower in AF13 cells compared to CTRLs across all cardiomyocyte subtypes (**Figure 6G**). Notably, AAV6 Cav1.3-C-terminus in the AF13 cells successfully rescued the uniform contraction of the monolayer in each cell subtype (**Figure 6G**).

**Figure 6:**
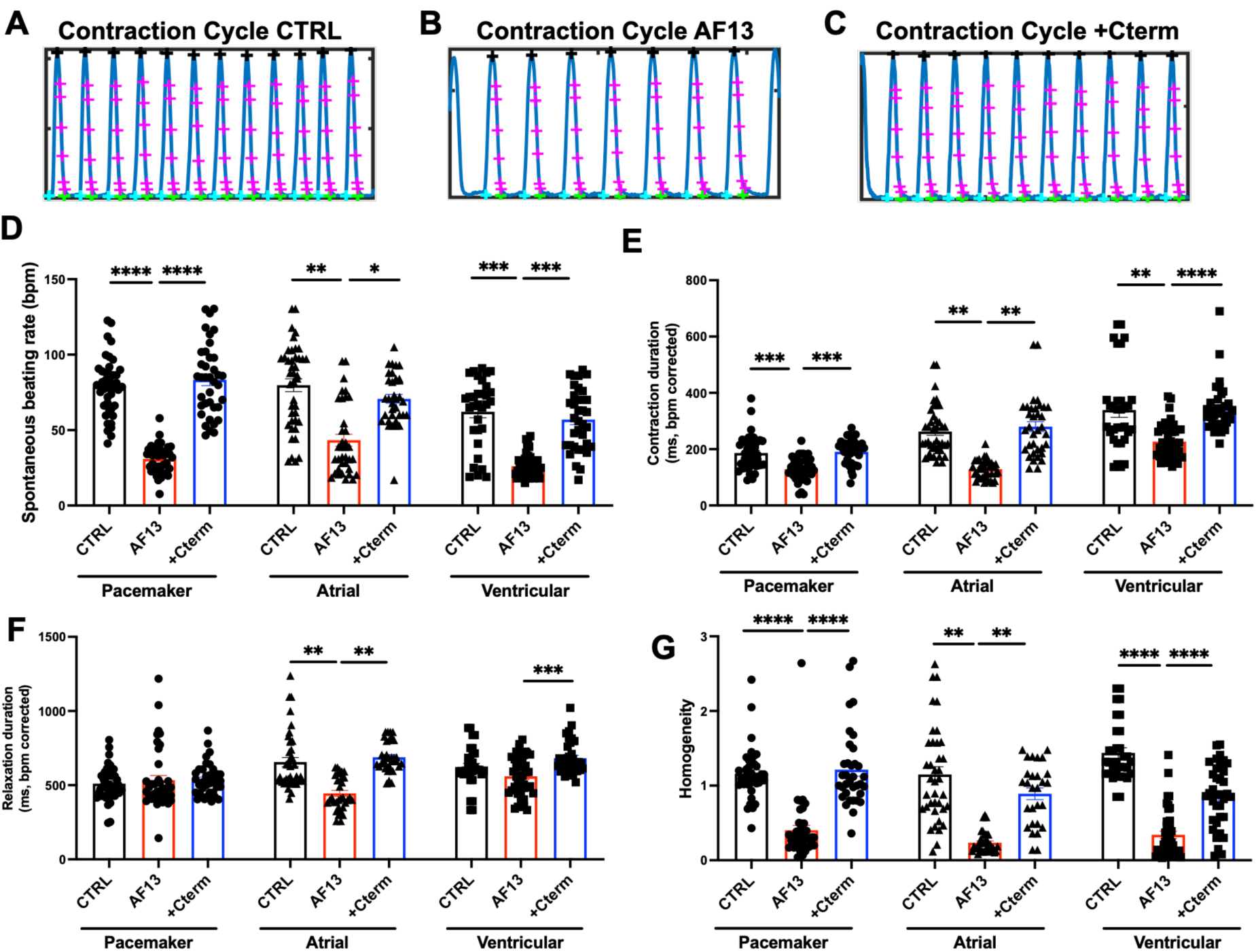
Evaluation of CTRL, AF13 and +Cterm spontaneous contractile function by video microscopy in cardiomyocytes subtypes. Contraction cycles from phase-contrast videos of CTRL **(A)**, AF13 **(B)** and after AAV6 Cav1.3-C-terminus treatment (+Cterm) **(C)** hiPSC-CMs monolayers obtained using custom made software analysis. **(D-G)** Contraction characteristics of hiPSC-PMs CTRL (n=43), AF13 (n=41) and +Cterm (n=38); hiPSC-aCMs CTRL (n=42), AF13 (n=36) and +Cterm (n=35); and hiPSC-vCMs CTRL (n=32), AF13 (n=43) and +Cterm (n=36). **(D)** Spontaneous beating rate, **(E)** Contraction and **(F)** Relaxation duration corrected by the bpm, **(G)** Homogeneity of the contraction. N = 3 independent differentiations. Error bars, s.e.m. Kruskal-Wallis Test. *p < 0.05, **p < 0.01, ***p < 0.001, ****p < 0.0001. CTRL: Control hiPSC-CMs; AF13: hiPSC-CMs carrying Cav1.3 variant; +Cterm: AF13 + AAV6 Cav1.3-C-terminus treatment.

The propagation of calcium influx in monolayer hiPSC-CMs monolayer was investigated by optical mapping recordings (**Figure 7**). Activation maps for hiPSC-PMs monolayer from the optical recordings (**Figure 7A**) showed that the propagation velocity (PV) was slower by 50% in AF13 than the CTRL and was normalized by the AAV6 Cav1.3-C-terminus treatment (**Figure 7B**). The calcium transient (CaT) characteristics were also modified, including a decrease in upstroke velocity and CaT decay time at 80% (CaT TD80) compared to CTRL, both of which were normalized after AAV6 Cav1.3-C-terminus treatment (**Figure 7C and D**). Activation maps of hiPSC-aCMs monolayer demonstrated also a 45% slower PV in AF13 compared to CTRL, which was rescued by the AAV6 Cav1.3-C-terminus treatment (**Figure 7E and F**). The CaT parameters were also modified, showing a decrease of upstroke velocity and CaT TD80 in AF13 compared to CTRL monolayer, which were normalized with the AAV6 Cav1.3-C-terminus treatment (**Figure 7G and H**). Finally, the activation maps of hiPSC-vCM monolayers showed a 58% slower PV in AF13 compared to CTRL but was not significantly normalized after the AAV6 Cav1.3-C-terminus treatment despite an upward reversal trend, as observed in other cardiomyocytes subtypes (**Figure 7I and J**). In addition, the CaT parameters showed a decrease of upstroke velocity in AF13 compared to CTRL, with a tendency to normalize this parameter with the AAV6 Cav1.3-C-terminus treatment but was not significant compared to AF13 (**Figure 7K**). The CaT TD80 was decreased in AF13 compared to CTRL and normalized after the AAV6 Cav1.3-C-terminus treatment (**Figure 7L**).

**Figure 7:**
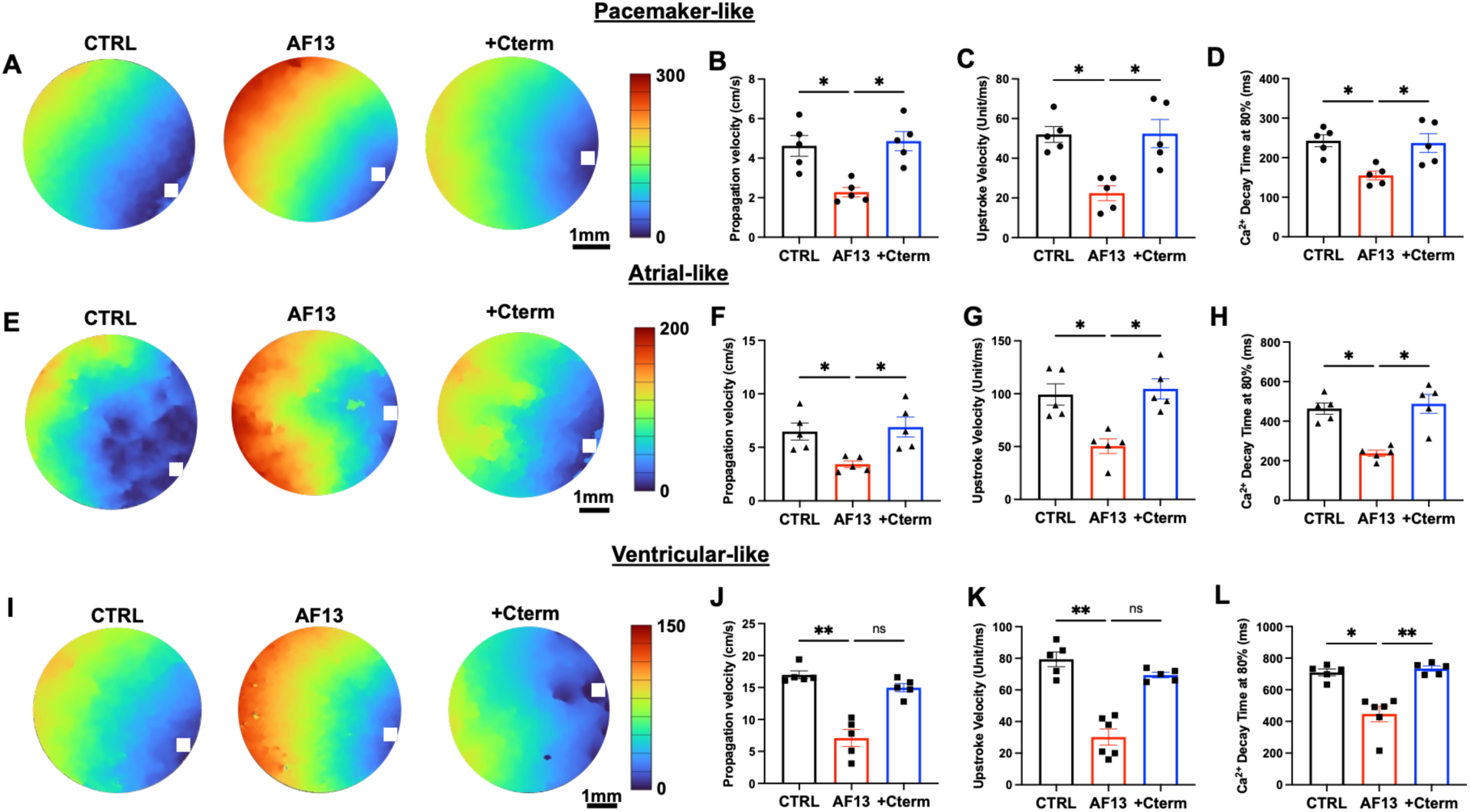
Evaluation of calcium conduction by optical mapping in CTRL, AF13 and +Cterm condition in cardiomyocyte subtype monolayers. **(A)** Representative activation map at a pacing of 1 Hz in hiPSC-PMs CTRL, AF13 and after AAV6 Cav1.3-C-terminus treatment (+Cterm); blue represents the early activation area and red the late activation area. Bar graphs summarizing **(B)** the calcium propagation velocity, **(C)** the upstroke velocity of the calcium transient and **(D)** the calcium decay time at 80% in hiPSC-PMs. **(E)** Representative activation map at a pacing of 1 Hz in hiPSC-aCMs CTRL, AF13 and +Cterm. Bar graphs summarizing **(F)** the calcium propagation velocity, **(G)** the upstroke velocity of the calcium transient and **(H)** the calcium decay time at 80% in hiPSC-aCMs. **(I)** Representative activation map at a pacing of 1 Hz in hiPSC-vCMs CTRL, AF13 and +Cterm. Bar graphs summarizing **(J)** the calcium conduction velocity, **(K)** the upstroke velocity of the calcium transient and **(L)** the calcium decay time at 80% in hiPSC-aCMs. N = 5 independent differentiations. White square show the localization of activation site. Error bars, s.e.m. Kruskal-Wallis Test. *p < 0.05, **p < 0.01. CTRL: Control hiPSC-CMs; AF13: hiPSC-CMs carrying Cav1.3 variant; +Cterm: AF13 + AAV6 Cav1.3-C-terminus treatment.

In addition to analyzing PV and CaT characteristics, we conducted an arrhythmia induction protocol (S1-S2 stimulation) to determine if the patient’s cells were more susceptible to arrhythmia. For this purpose, triggered arrythmia was defined as a sustained VT lasting longer than 1 minute. In CTRL monolayer, the S1-S2 stimulation triggered arrhythmia in only 20% of monolayers. In most instances, the S1-S2 protocol was followed by spontaneous activity at 1Hz with 200 ms, 400 ms or 500 ms S2 for hiPSC-PMs, -aCMs or -vCMs respectively (**Figure 8A and B**). Notably, AF13 monolayer exhibited a much higher susceptibility, with arrhythmias being triggered in 60% of monolayers. These arrhythmias appeared to be a form of spiral wave reentry (**Figure 8C and D**). Upon AAV6 Cav1.3-C-terminus treatment, the AF13 monolayer showed a significant reduction in arrhythmia susceptibility, with arrhythmias triggered in only 30% of monolayers. In most instances, the S1-S2 protocol, like the CTRL condition, was followed by normal spontaneous activity (**Figure 8E and F**). These results collectively demonstrate that the *de novo CACNA1D* variant impairs the contractile function, calcium wave PV and increase arrhythmia susceptibility in hiPSC-CMs monolayer. More importantly, the AAV6 Cav1.3-C-terminus treatment was able to restore normal contraction and calcium wave properties and reduce arrhythmic events.

**Figure 8:**
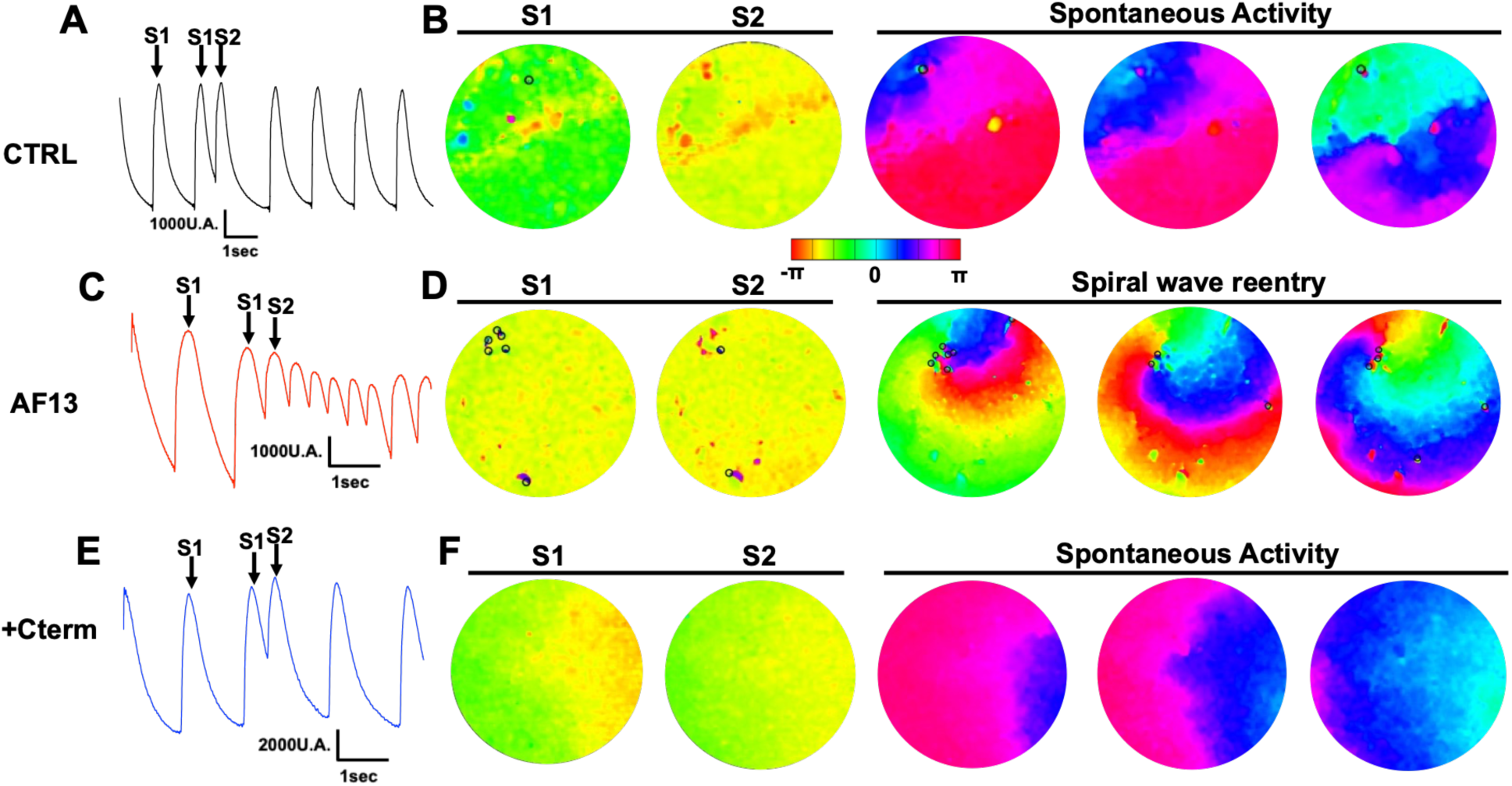
Evaluation of arrhythmia inducibility in hiPSC-CMs monolayer. **(A)** Raw traces of calcium transient response to the S1-S2 stimuli and **(B)** the corresponding representative phase maps during and after the S1-S2 protocol in CTRL hiPSC-CM monolayer. **(C)** Raw traces of calcium transient response to the S1-S2 stimuli and **(D)** the corresponding representative phase maps during and after the S1-S2 protocol in AF13 hiPSC-CMs monolayer. **(E)** Raw traces of calcium transient response at the S1-S2 stimuli and **(F)** the corresponding representative phase maps during and after the S1-S2 protocol in +Cterm hiPSC-CMs monolayer. N = 5 independent differentiations.

## Discussion

We investigated the consequences of the expression of a *de novo* variant of the *CACNA1D* gene from a pediatric patient on the function of chamber-specific hiPSC-CMs. This gene encodes the L-type calcium channel Cav1.3, which is a crucial regulator of calcium entry in cardiomyocytes, particularly in controlling the automaticity of pacemaker cells and the propagation of electrical impulses in the AVN^9,36^. The patient with this variant presented with sinus bradycardia, conduction abnormalities, and episodes of ventricular tachycardia, phenotypes that we were able to reproduce in the same patient-specific hiPSCs but not control sex-and age-matched. We demonstrated a loss of function in Cav1.3, evidenced by a decrease in gene expression and a disruption of the membrane localization of the Cav1.3 channel. Furthermore, hiPSC-CMs carrying the variant, across all cells subtypes (i.e., pacemaker, atrial and ventricular cells), showed a decrease in ICaL densities along with other electrophysiological disturbances, including a reduced APD, spontaneous beating rate, calcium propagation wave and increased susceptibility to arrhythmia induction. These findings provide electrophysiological basis for the clinical electrical phenotype observed in the patient. We then tested a gene therapy approach to restore the loss of Cav1.3 function by upregulating the expression of Cav1.3. This novel approach involves using an AAV6 vector to deliver the Cav1.3-C-terminus to hiPSC-CMs. Our study expands the understanding of Cav1.3’s role in cardiac arrhythmias and illustrates the promise of precision medicine in developing targeted treatments for specific genetic defects.

The LTCCs are a key regulators of calcium balance in cardiomyocytes, allowing rapid entry of calcium into the cytosol, which activates myofilaments and produces contraction^37^. In HF, for example, dysregulation of intracellular calcium handling is a primary characteristic causing contractile dysfunction and arrhythmias^38^. In the heart, LTCCs are composed of the Cav1.2, ubiquitously expressed in all heart chambers, and the developmentally regulated Cav1.3, exclusively expressed in the atria, SAN, and AVN in adult hearts^39^. In the SAN, Cav1.3 regulates the automaticity of the pacemaker cells^40,41^. Previous studies show that genetic deletion of *CACNA1D* in mice,^14^ as well as the expression of *CACNA1D* variants in patients^15,42^ can caused SANDD, sinus bradycardia, and AVB, consistent with region-specific expression. Here, we describe a newly identified, *de novo* variant of *CACNA1D* (c.3786G>T) found in a 13-year-old pediatric patient with a history of VT episodes dating back to age seven. Genetic screening of both the patient and the mother identified two additional variants (*SCN5A* (c.2618C>G and *DSP* c.1582C>G). The *SCN5A* gene, which encodes the Nav1.5 channel, is a well-known risk factor for Brugada syndrome and long QT syndrome, while the *DSP* gene, responsible for the expression of desmoplakin, a key desmosomal protein, implicated in arrhythmogenic cardiomyopathy (ACM)^43,44^. As the patient’s mother is asymptomatic but presents with Brugada-like ECG and had no history of arrhythmias or structural heart abnormalities, her inherited genetic variants (*SCN5A* and *DSP*) likely did not contribute to the AF13 patient’s clinical presentation but may have provided the needed substrate for *CACNA1D* to unmask the arrhythmogenic phenotype.

This underscores the importance of our study, which aims to thoroughly explore the effects of the *CACNA1D* variant on the electrophysiological properties of cardiomyocytes and its contribution to arrhythmias.

To decipher the role of this variant, we used hiPSCs derived from the patient. The hiPSC model is a valuable tool for studying channelopathies specific to the patient^45^. In this study, we have differentiated the hiPSCs into 3 cardiomyocytes subtypes (pacemaker, atrial and ventricular) and were able to reproduce and investigate the patient’s phenotypes. We found that *CACNA1D* gene expression was reduced in the patient’s hiPSC-CMs; the latter was accompanied by a reduced abundance of Cav1.3 protein localization at the sarcolemma when compared to CTRL (Figure 2, Supplemental Figures 1 and 2). In CTRL hiPSC-CMs, Cav1.3 is properly localized around the nucleus and the cell membrane^46^. However, in the patient’s cells, the protein is absent from these locations, confirmed by the immunofluorescence experiments. This lack of proper localization suggests that the mutated Cav1.3 protein is likely misfolded and, therefore, unable to be transported to its correct destination. This is a common encounter with misfolded proteins, which often display endoplasmic reticulum (ER)-retention signals, causing their retention in the ER and their subsequently degradation by the proteasome. This process, known as ER-associated degradation (ERAD), is a quality control mechanism that prevents misassembled ion channels from reaching the cell membrane, ultimately disrupting the entire trafficking pathway. An other explanation of this lack of protein localization could be possible with a decrease of RNA translation into proteins^47^. This reduction of Cav1.3 expression and localization were linked to a reduction of ICaL densities, as observed in patch-clamp experiments. No Cav1.3 currents kinetics were observed, the decrease in current densities can be explained by a decrease of functional Cav1.3 channels.

We also identified several alterations in AP parameters in the patient’s hiPSC-CMs, including reduced APD, amplitude, and upstroke velocity. The latter two may be associated with a decrease in INa density. Indeed, the *SCN5A* variant introducing a premature stop codon is highly pathogenic. Moreover, the *DSP* variant may further compromise sodium channel anchoring at the intercalated disc, thereby contributing to the reduced INa observed in patch-clamp recordings^48^. A reduction in both sodium and calcium entry is known to alter these AP electrophysiological properties^49^. Specifically, a shorter APD promotes arrhythmias by decreasing the refractory period. This makes the cardiac tissue more susceptible to re-excitation, which can lead to arrhythmic events^50,51^. The shortened AP was associated with an increase in potassium channel expression observed by RNA-seq. Finally, a decrease in spontaneous beating rate and calcium wave propagation was observed in patient’s hiPSC-CMs; the latter can be attributed to the Cav1.3 loss of function, in the absence of changes in *HCN* transcript abundance^40^. Furthermore, the Cav1.3 channel plays a critical role in the electrical conduction of the AP in the SAN, and AVN^8,52^. We therefore speculate that the reduction in ICaL density was a major contributor to the slowing of cardiac impulse propagation in the patient’s heart^53^. Interestingly, hiPSC-CMs from the patient were more prone to developing arrhythmias after an S1-S2 induction protocol. Based on previous observations, a decrease in ICaL disturbs the electrophysiological properties of cardiomyocytes, leading to a shorter refractory period, reduced calcium wave propagation, and a slower beating rate. All these factors are pro-arrhythmic and may contribute to re-entrant arrhythmias^54^. We observed Cav1.3 expression in the ventricular cardiomyocytes, which is typically absent in adult hearts. This expression may persist in these hiPSC-vCMs, consistent with the pediatric status of the patient. The first part of this present study seeks to explain the phenotypes observed between the mother and the daughter. Specifically, some inherited channelopathies exhibit incomplete or reduced penetrance, such as ACM^55^ and Brugada syndrome^56^, where individuals who inherit a predisposing gene variant do not develop the disease. The mother, who presents a Brugada-like ECG, does not exhibit any arrhythmias or structural fibro-fatty replacement of the right ventricle, would otherwise be suggested by the *DSP* variant. This statement support the “multihit-theory” where a single channelopathy is not able in some cases to induce symptoms, and rarely even the related clinical phenotype^57,58^. This theory is supported by our results, which show a reduction in calcium entry observed in the daughter’s hiPSCs. This suggests that the addition of *CACNA1D* variant could be a key factor triggering the daughter’s phenotype. It is important to note that calcium handling disturbances are implicated in, and contribute to, both Brugada syndrome^59^ and ACM^60^. In ACM, the arrhythmias and electrophysiological remodeling are the initial and direct pathological mechanism; they are not a consequence of subsequent tissue remodeling^30,61^. This explains the absence of ACM evidence in the daughter via cardiac MRI.

To attempt a novel therapeutic approach into rescuing this observed Cav1.3 loss of function in hiPSC-CMs, we leveraged Cav1.3-C-terminus’s unique intrinsic property of acting as a transcriptional factor which upregulates the expression of its own gene^19,20^. We previously demonstrated in a mouse model of HF that added expression of the C-terminus fragment upregulated Cav1.3 expression and increased the LTCCs currents^20^. In this present study, the Cav1.3-C-terminus was integrated into an AAV6 vector for delivery into the patient’s hiPSC-CMs, as previously described^21^. The treatment led to the upregulation and re-localization of Cav1.3, which normalized the cells’ electrophysiological and mechanical properties. Furthermore, this treatment not only restored a normal expression of Cav1.3, but also other genes (such as potassium channel genes) which contribute to a normalization of function. Interestingly, the AAV6 Cav1.3-C-terminus treatment improved the sarcomere organization, as assessed by immunofluorescence for the α-actinin marker, likely due to the normalization of calcium entry to the cell^62,63^. It is important to add that the Cav1.3-C-terminus treatment did not only affect the expression of its own gene, but also the overall transcriptome. In fact, calcium act like a second messenger for the transcriptional regulation of many genes^62^. Disturbances of cellular calcium handling affect not only excitation-contraction coupling, but also calcium-dependent gene expression. The normalization of calcium entry by the Cav1.3-C-terminus has a dual effect; it restores functional contraction and normalize the expression of genes. On the other hand, the Cav1.3-C-terminus treatment could also affect the expression of gene detrimental to the cardiomyocyte integrity, but our results show a decrease in the risk of arrhythmogenesis in the treated hiPSC-CMs. Taken together, this gene therapy approach can reduce arrhythmia risk by restoring the calcium homeostasis back to a normal state. These results are promising for future treatments of cardiac arrhythmias that involve loss of function in LTCCs.

In the present study, we describe a multi-hit theory based on the investigation of a *de novo CACNA1D* variant, which accounts for the daughter’s phenotypes that are not present in the mother. This work could be enhanced by future studies generating isogenic controls for each variant of interest (*CACNA1D* (c.3786G>T), *SCN5A* (c.2618C>G), and *DSP* (c.1582C>G)). This methodology would reinforce our multi-hit theory by allowing to correct and confirm the pathogenicity of the respective variants. However, the genomes of selected clones may differ. These differences can arise from unintended edits by the Cas9 enzyme, known as off-target effects. More significantly, genomic differences can stem from the natural accumulation of variants in the cells of the parental iPSC lines. While these latter variants are present in only a small percentage of cells in the line, they can be disregarded as a potential source of phenotypic variation^64,65^. This problem could lead to bias and misinterpretation. Furthermore, this study is based on a single hiPSC line. It would be valuable to confirm our data on the Cav1.3-C-terminus treatment in other hiPSC lines carrying distinct LTCC loss of function variants, or in model of hiPSCs that reproduce HF.

In summary, our study used hiPSC-CMs to uncover molecular mechanisms in human subject and develop precision therapeutic strategies. We determined that the additional *CACNA1D* c.3786G>T variant in the daughter causes a loss of function in the Cav1.3 channel, which explains the arrhythmias seen in the patient. Building on this proof of concept, we developed a promising gene therapy approach that could be used for cardiac diseases caused by loss of calcium influx. The Cav1.3-C-terminus, a key component that can promote the expression of its own gene, was able to restore a normal calcium balance and reduce the risk of arrhythmias. This work is a crucial step toward future clinical trials for new cardiovascular therapies that aim to prevent calcium handling disturbances and related cardiac dysfunction.

## Data availability

The data supporting this article and other findings are available within the manuscript, figures, supplementary figures and from the corresponding author upon request.

## Acknowledgements

We thank the platform Neurophotonics from Québec, Canada that provided the AAV6 and plasmid containing the Cav1.3 C-terminus construct. CTRL hiPSCs cell line was obtained as a generous gift from the Stanford Cardiovascular Institute (SCVI) Biobank.

We would like to acknowledge our fundings from Biomedical Laboratory Research & Development Service of United States (U.S.) Department of Veterans Affairs, Office of Research and Development, Merit Review grant I01 BX002137 to MB; National Heart, Lung, and Blood Institute 1R01HL164415 to MB; US Department of Defense award number W81XWH-21-1-0424 to MB. This work was also supported by the American Heart Association Postdoctoral Fellowship (https://doi.org/10.58275/AHA.25POST1361676.pc.gr.227353) to J-BR.

## Author contributions

J-B.R., Y.S. and M.B designed the experiments, contributed to the discussion, and wrote the manuscript. J-B.R., Y.S., H.DA., M.C., V.K.M.G collected and analyzed the data. R.T., W.C., YS. Q. and F.C. performed the clinical approaches and the sampling blood collection from patient. J-B.R., M.D., M.C. and M.B. contributed to the discussion and reviewed/re-edited the manuscript.

