## Supplemental Figure for "Electrophysiological abnormalities associated with a *CACNA1D* variant are rescued by AAV6-Cav1.3-C-terminus gene therapy in patient-iPSC-CMs"

#### Supplemental Figure 1

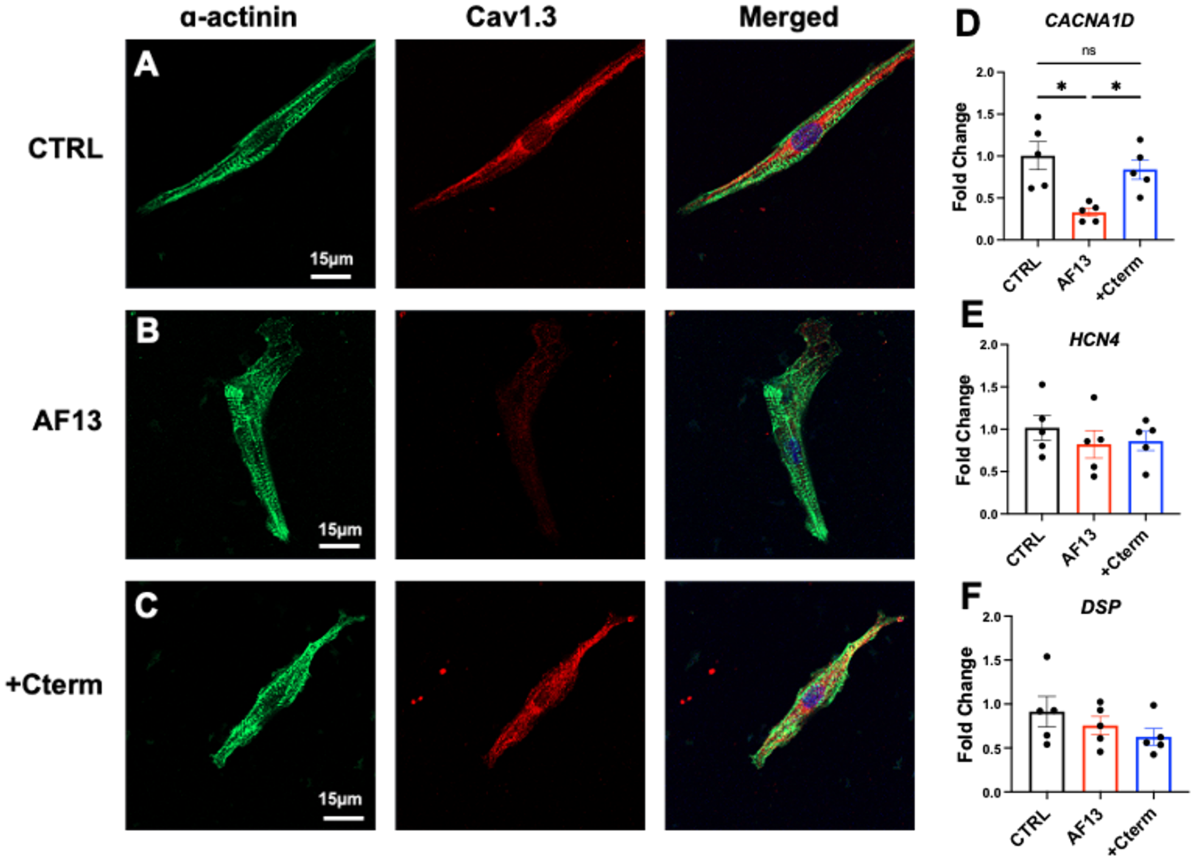


**Supplemental Figure 1: Molecular characterization of Cav1.3 variant in pacemaker-like hiPSC-PMs.** Immunostaining of α-actinin (green), Cav1.3 (red) and nuclei (DAPI, cyan) were acquired in CTRL **(A)**, AF13 **(B)** and after treatment with AAV6 Cav1.3-C-terminus **(C)**. RT-qPCR analysis of other genes implicated in cellular excitability (*CACNA1D* and *HCN4* **(D and E)**) and in cardiomyocytes adhesion (*DSP* **(F)**). Results are shown as mean±s.e.m. * p<0.05, ns: not statically significant (Kruskal-Wallis test). CTRL: Control hiPSC-CMs; AF13: hiPSC-CMs carrying Cav1.3 variant; +Cterm: AF13 + AAV6 Cav1.3-C-terminus treatment.

#### Supplemental Figure 2

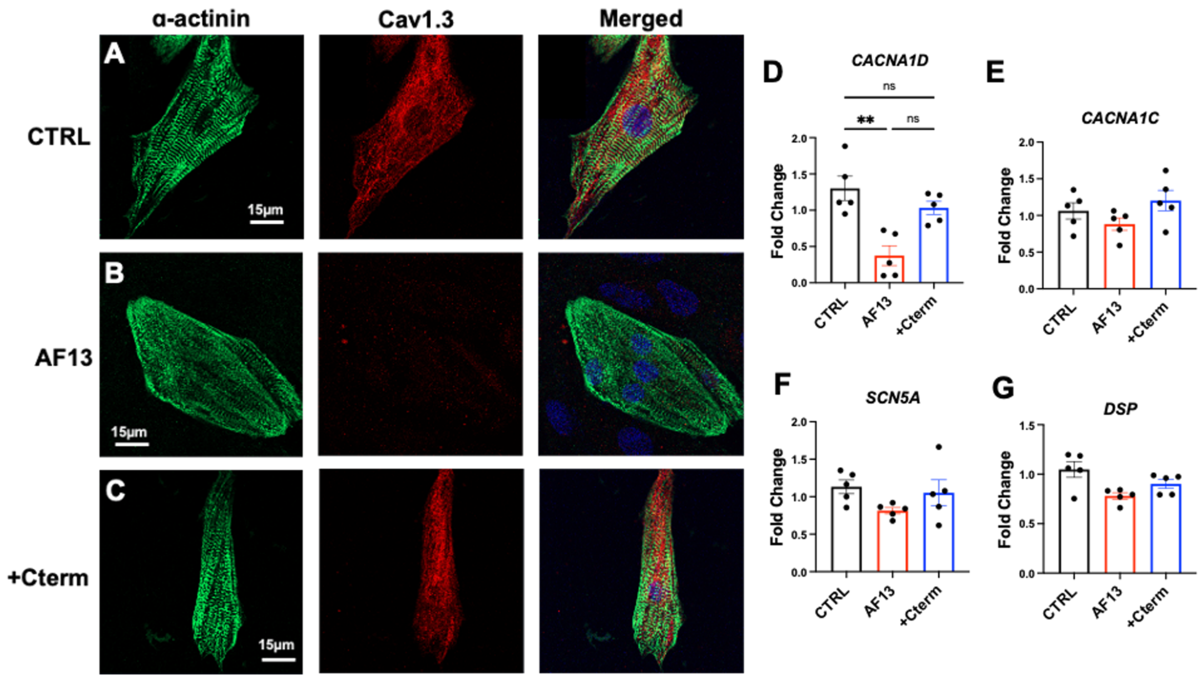


**Supplemental Figure 2: Molecular characterization of Cav1.3 variant in ventricular-like hiPSC-vCMs.** Immunostaining of α-actinin (green), Cav1.3 (red) and nuclei (DAPI, cyan) were acquired in CTRL **(A)**, AF13 **(B)** and after treatment with AAV6 Cav1.3-C-terminus **(C)**. RT-qPCR analysis of other genes implicated in cellular excitability (*CACNA1D*, *CACNA1C* and *SCN5A* **(D-F)**), cell adhesion (*DSP* **(G)**). Results are shown with the mean±s.e.m. * $p<0.05$, ns not statically significant (Kruskal-Wallis test). CTRL: Control hiPSC-CMs; AF13: hiPSC-CMs carrying Cav1.3 variant; +Cterm: AF13 + AAV6 Cav1.3-C-terminus treatment.

### Supplemental Figure 3

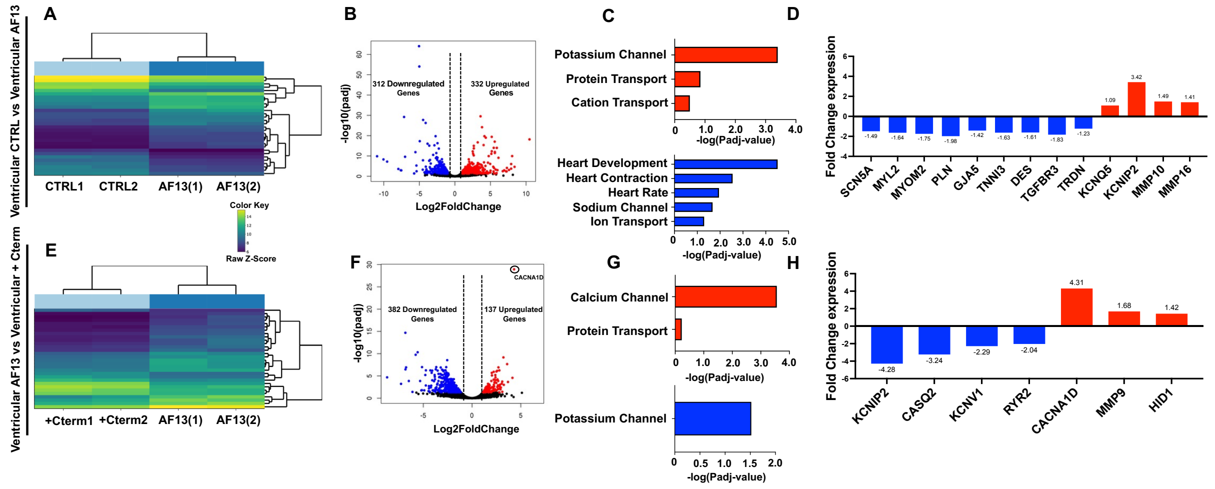


**Supplemental Figure 3: Transcriptomic analysis of human induced pluripotent stem cell-derived ventricular-like cardiomyocytes (hiPSC-vCMs). (A)** Heat map plot of the top 30 DEGs between CTRL and AF13. **(B)** Volcano plot of downregulated (blue) and upregulated (red) genes in CTRL and AF13. **(C)** Gene ontology biological analysis of DEGs in CTRL and AF13. **(D)** Fold change in expression of key genes. **(E)** Heat map plot of the top 30 DEGs between AF13 and after AAV6 Cav1.3-C-terminus treatment (+Cterm). **(F)** Volcano plot of downregulated (blue) and upregulated (red) genes in AF13 and +Cterm. (G) Gene ontology biological analysis of DEGs in AF13 and +Cterm. **(D)** Fold change in expression of key genes. The Wald test was used to generate p-values and log2 fold changes. Genes with an adjusted p-value < 0.05 and absolute log2 fold change > 1 were considered as differentially expressed genes

#### Supplemental Figure 4

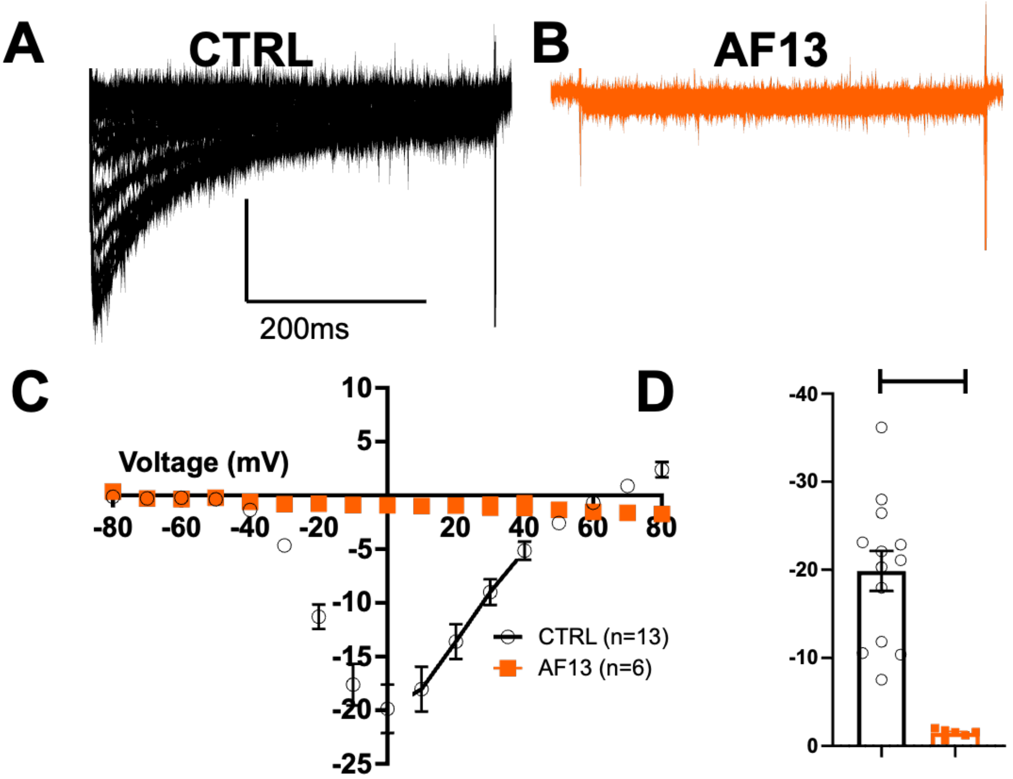


**Supplemental Figure 4: Evaluation of Cav1.3 variant in tsA201 cells.** Representative calcium currents traces recorded in **(A)** CTRL (WT + $Ca_V\alpha2\delta1$ WT + $Ca_V\beta3$ WT (GFP)) (black; n=13); **(B)** AF13 ($Ca_V1.3$ M1262I + $Ca_V\alpha2\delta1$ WT + $Ca_V\beta3$ WT) (orange; n=7) in tsA201 cells. **(C)** Normalized Current/Voltage curves. **(D)** Peak current density at 0 mV. Statistical comparisons were performed by Student's t tests. Bars indicate s.e.m. ****$p < 0.0001$

#### Supplemental Figure 5

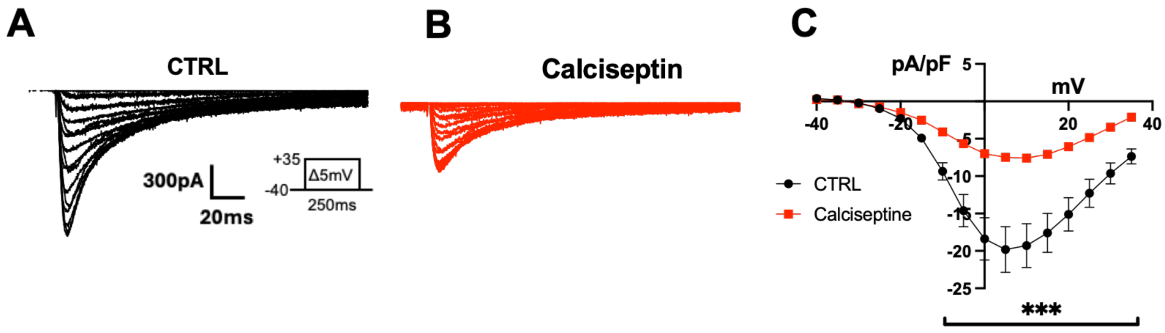


**Supplemental Figure 5: Evaluation of Cav1.3 current by the calciseptine.** Representative calcium current recorded in voltage-clamp mode in **(A)** CTRL (black; n=8); **(B)** after calciseptine (1 μM) application (red; n=8) in hiPSC-CMs. **(D)** Normalized Current/Voltage curves in hiPSC-vCMs. N = 2 independent differentiations. Error bars, s.e.m. Kruskal-Wallis Test. ***p < 0.001.

#### Supplemental Figure 6

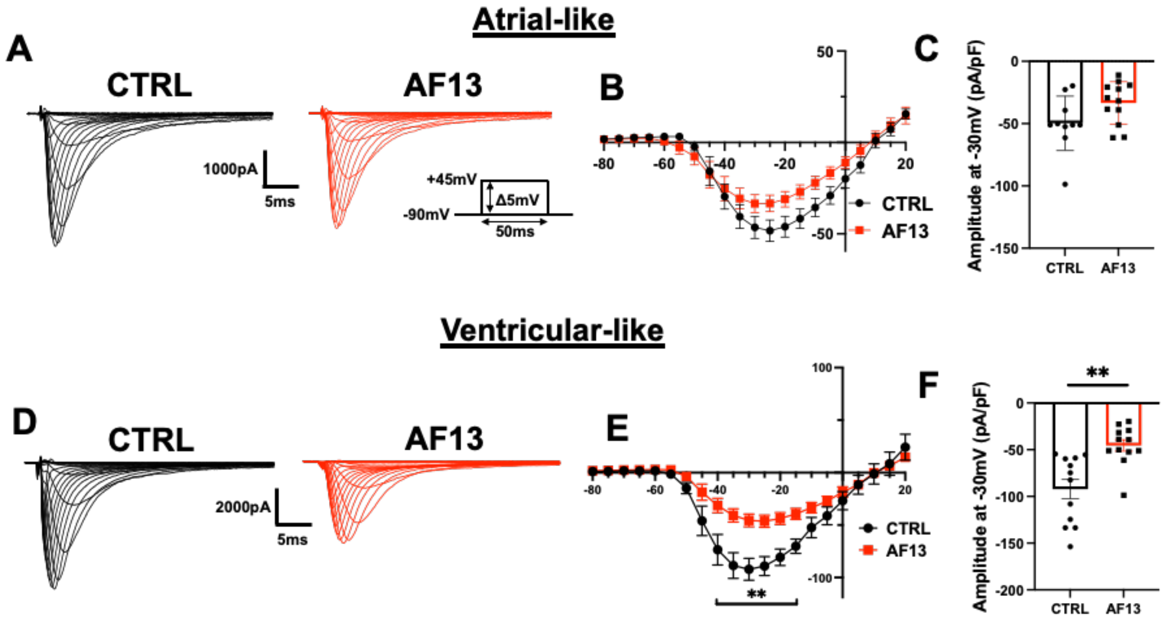


**Supplemental Figure 6:** Biophysical properties of sodium channels in hiPSC atrial- and ventricular-like cells. Representative sodium current recorded in voltage-clamp mode in **(A)** CTRL (black; n=10) and AF13 (red; n=12) in hiPSC-aCMs. **(B)** Normalized Current/Voltage curves with **(C)** the corresponding current amplitude measurements at -30mV. Representative sodium current in **(D)** CTRL (black; n=12) and AF13 (red; n=12) in hiPSC-vCMs. **(E)** Normalized Current/Voltage curves with **(F)** the corresponding current amplitude measurements at -30mV. N = 3 independent differentiations. Error bars, s.e.m. A Mann-Whitney test was performed. $^{**}p < 0.01$. CTRL: Control hiPSC-CMs; AF13: hiPSC-CMs carrying Cav1.3 variant.
